# Macronutrient Composition and Genetic Background Determine the Response to a Ketogenic Diet

**DOI:** 10.64898/2026.04.23.720368

**Authors:** Zhaoyue Zhang, Alexandre Moura-Assis, Shanshan Liu, Alon Millet, Jordan Shaked, Divya Rajan, Hanan Alwaseem, Michael Isay-Del Viscio, Henrik Molina, Kivanc Birsoy, Jeffrey Friedman

## Abstract

While standard high fat diets cause hyperphagia and obesity in mice, high fat-low carbohydrate ketogenic diets (KDs) reduce food intake and body weight. Because the basis for this difference is still unclear, we systematically altered the macronutrient content of a standard KD and found that feeding C57BL/6J (B6J) mice a KD with 5% protein resulted in hypophagia, weight loss, and hypoglycemia, whereas the same diet with 10% protein led to increased adiposity and glucose intolerance. However, these effects were strain-dependent as C57BL/6NJ (B6NJ) weighed similar amounts on the two diets leading us to investigate the molecular mechanisms. When fed the KD-5% diet, B6J but not B6NJ mice showed increased levels of two anorexigenic factors, GDF15 and LCN2, and loss of function of either blunted the weight loss of B6J mice fed the diet. B6J mice harbor mutations in Nnt (Nicotinamide nucleotide transhydrogenase) and Nlrp12 (NLR family pyrin domain containing 12), both of which are wildtype in B6NJ mice. B6J mice fed the KD-5% diet showed the RNA signature of oxidative and integrated stress responses (ISR) and restoring NNT function in liver reduced the levels of GDF15. RNA-seq also revealed that B6J but not B6NJ mice had the RNA signature for hepatic inflammation and a knockout of Nlrp12 led B6NJ mice to lose weight on the KD-5% diet with increased levels of LCN2. Suppression of oxidative stress with N-acetylcysteine (NAC) reduced expression of both GDF15 and LCN2 and prevented the weight loss associated with the KD-5% protein diet in B6J mice, whereas inhibition of the integrated stress response with ISRIB only attenuated the GDF15 axis. Collectively, these findings explain why B6J mice lose weight on a ketogenic diet and reveal a critical interplay between macronutrient composition and genetic background leading to increased levels of GDF15 and LCN2 to induce hypophagia. Finally, these data suggest that the response to different diets among humans might be similarly variable based on genetic variation and macronutrient composition, suggesting the possible need for personalized dietary interventions.

## Introduction

Genetic and environmental factors both contribute to the development of obesity in humans and mice^1^. In humans, twin studies, adoption studies, studies of familial aggregation as well as molecular genetic analyses suggest a substantial genetic component while the secular trend toward increased obesity over time suggests that environmental factors also contribute^1,2^. In mice, several Mendelian loci cause obesity^3^ and while some mouse strains fed a high fat diet develop severe obesity others do not^3^. For instance, similar to many humans consuming “Western” diets, C57BL/6J (B6J) mice fed high-fat diets (HFD) develop leptin resistance and obesity while other strains such as A/J and CAST/EiJ mice remain lean^4–6^. Thus, the aggregate data in both species suggests that obesity is caused by a gene-environment interaction with the macronutrient composition of a diet influencing the outcome.

Ketogenic diets (KDs), characterized by high fat and minimal carbohydrate content, exert profound metabolic effects in both humans and rodents^7,8^. Indeed, ketogenic diets (KDs) with a composition of 90% or more of fat, moderate protein, and minimal carbohydrates, have been used to treat seizures, obesity and other disorders^9–13^. However, while KDs have been widely used in multiple clinical settings to these disorders^14^, the clinical and pre-clinical effects on food intake, body weight and glucose homeostasis have been inconsistent^15–18^ and the basis for this variability has proven elusive. While the anorexigenic factor GDF15 has been reported to contribute to KD-induced weight loss, its blood levels have been similarly variable, with some, but not all, studies showing elevated levels raising the possibility that there are additional modulatory factors^19,20^. In addition, the mechanism leading to activation of GDF15 in animals fed a KD is not fully understood.

To address this, we systematically varied the macronutrient composition of a KD in B6J mice. The aggregate studies revealed that feeding C57BL/6J mice a KD with 5% protein reduces food intake and body weight while improving glucose tolerance, whereas the same diet with 10% protein (KD-10%P) increases adiposity and causes glucose intolerance. Studies of glucose metabolism indicated that the differential effects of the 5% protein KD vs. the 10% protein KD were largely substrate driven.

Surprisingly, we also found that the differential effects of the 5% protein KD vs. the 10% protein KD on food intake and body weight were strain dependent with C57BL/6NJ (B6NJ) showing no difference. B6J mice carry mutations in both the Nnt gene, which regulates mitochondrial redox balance, and Nlrp12, an anti-inflammatory regulator, raising the possibility that these mutations might contribute to weight loss on a KD-5%P^21^. We found that in animals lacking functional NNT, KD-5%P feeding triggers hepatic stress responses leading to elevated GDF15 levels and weight loss. In parallel, dysfunction of NLRP12 was associated with increased inflammatory responses and elevated levels of the anorexigenic factor LCN2^22^, further amplifying hypophagia. Both responses and the associated weight loss are blunted by treatment with NAC which reduces oxidative stress. In aggregate, these data suggest that the variable effects of ketogenic diets are likely a result of differences in both genetic background and macronutrient composition which shape metabolic outcomes through coordinated activation of stress and immune pathways.

## Results

### Protein Content Dictates the Effects of a KD in B6J Mice

Ketogenic diets (KDs) are composed of high levels of fat, moderate protein, and little or no carbohydrate^13^. However, the composition of KDs (including those from Bio-Serv (F3666) and Envigo (#96355)) often vary with differences in protein and fat composition, potentially leading to variable outcomes^9,19,23–25^. To investigate how macronutrient composition affects metabolic outcomes in B6J mice, we varied the macronutrients in a series of KDs and observed their effects on 8-week-old B6J male mice over a 3 to 4-week period.

We began by adjusting the fat composition in KDs with 10% protein and no carbohydrate and found that feeding C57BL/6J mice three different diets rich in poly-unsaturated fat (PUFA), mono-unsaturated fat (MUFA) or saturated fat (SAFA) failed to elicit differential effects on energy balance with food intake and fat mass increasing in all three cases relative to chow-fed controls (Figure 1a). Similarly, glucose tolerance tests showed that all three KDs led to a significantly increased area under the curve indicative of impaired glucose tolerance relative to the chow-fed group (Figure 1b). Notably, we did not observe any differences in fasted glucose or ketone levels among the 3 KD groups (Supplementary Figure 1a, b). These results suggest that fat composition minimally influenced the KD outcomes in B6J mice, and that high fat diets containing 10% protein promote obesity rather than weight loss in B6J mice.

**Figure 1.**
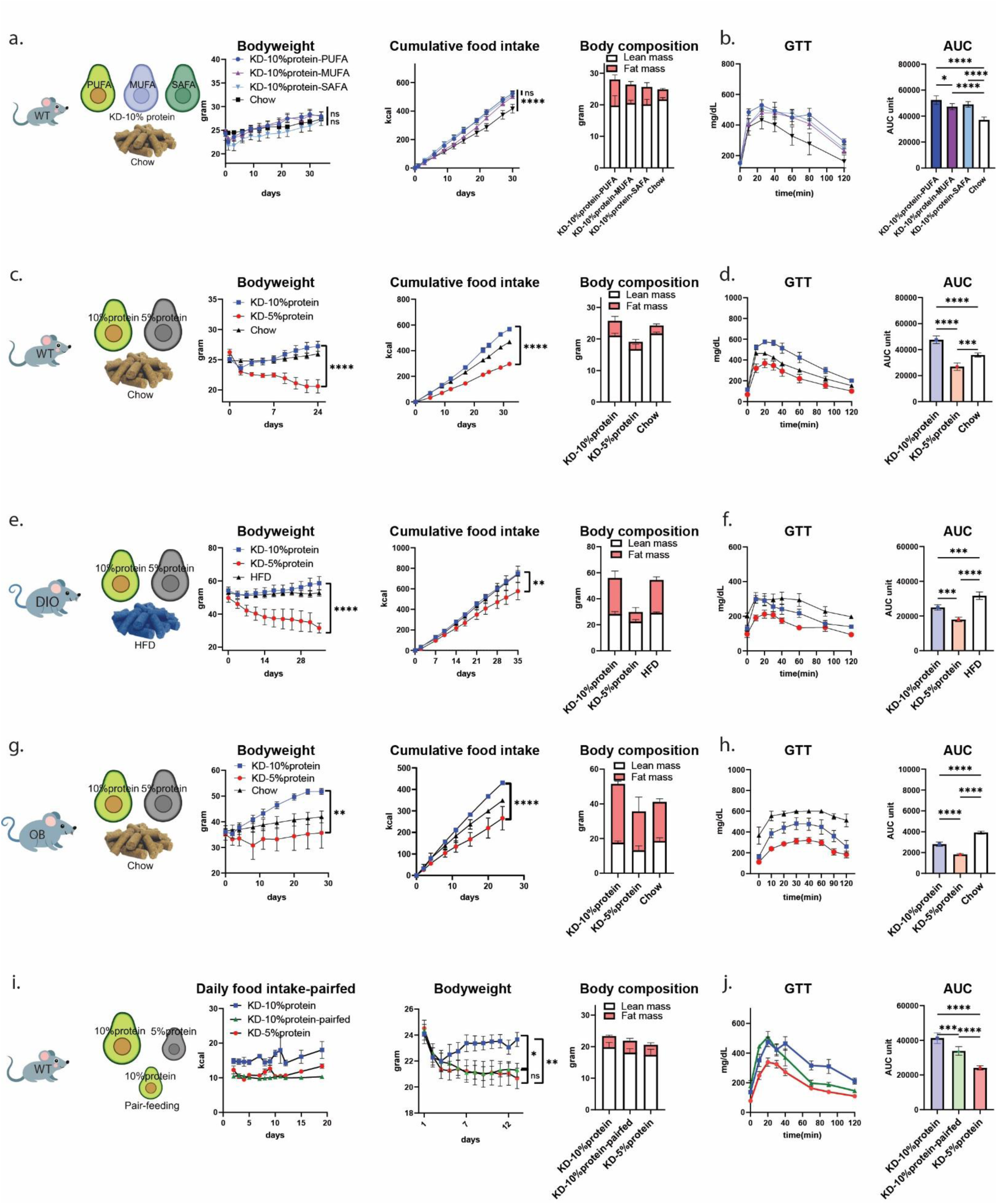
Protein Content Dictates KD Efficacy in B6J mice. a. The schematic of mouse strain and diets resulting in body weight, cumulative calories consumed, and body composition of WT B6J mice fed with chow, KD-10%P rich in PUFA, MUFA and SAFA after 4-week diet challenge. b. Glucose tolerance test (2mg/gram body weight) and calculated area under the curve of WT B6J mice fed with chow, KD-10%P rich in PUFA, MUFA and SAFA. c, d. As in a, b, for WT B6J mice fed with chow, KD-10%P, KD-5%P rich in PUFA. e, f. As in a, b, for DIO mice (Control diet is HFD; GTT is at the rate of 0.5 mg/gram body weight). g, h. As in a, b, for OB mice (GTT is at the rate of 0.5 mg/gram body weight). i. The schematic of pair-feed experiments and the result in body weight, daily calories consumed, and body composition of WT B6J mice fed with KD-5%P, KD-10%P and pair-fed KD-10%P to the amount of KD-5%P after 3-week diet challenge. j. Glucose tolerance test (2mg/gram body weight) and calculated area under the curve of WT B6J mice fed with KD-5%P, KD-10%P and pair-fed KD-10%P to the amount of KD-5%P. (n = 5 mice for all WT test, n=4 for all DIO test, n=4 for OB on KD and n=3 for OB on chow. Data are presented as mean ± s.e.m. NS, not significant; *P < 0.05; **P < 0.01; ***P < 0.001; ****P < 0.0001.)

We next varied protein levels in diets with the same fat composition of 52.8% PUFA, 21.4% MUFA and 25.8% SAFA. In contrast to the effect of diets with different lipid content, B6J mice fed a 5% protein KD (KD-5%P) exhibited a markedly reduced food intake, decreased body weight, and improved glucose tolerance compared to those fed the same diet with 10% protein (KD-10%P) (Figure 1c, d). After four weeks on KD-5%P, the B6J mice showed a ∼40% decrease in food intake (Figure 1c) (Average cumulative food intake for KD-10%P is 567.64±28.91kcal, KD-5%P is 295.99±27.11kcal, and chow is 467.58±28.16kcal. With the 2-way ANOVA test, the adjusted p value for KD-10%P vs. KD-5%P is <0.0001, for KD-10%P vs. chow is <0.0001, for KD-5%P vs. chow is<0.0001.), lost ∼ 30% of body weight (Figure 1c) (Average final body weight for KD-10%P is 27.21±1.88g, KD-5%P is 20.58±2.47g, and chow is 25.96±0.91g. With the 2-way ANOVA test, the adjusted p value for KD-10%P vs. KD-5%P is <0.0001, for KD-10%P vs. chow is 0.334, for KD-5%P vs. chow is<0.0001), with significant reductions in both lean and fat mass compared to chow-fed controls (Figure 1c). (Average lean mass for KD-10%P is 21.05±0.77g fat mass is 4.65±1.44g, KD-5%P is lean=16.76±1.58g fat=2.31±0.73g, and chow is lean=21.72±0.44g fat=2.55±0.52g). The KD-5%P group also had a decreased area under the curve during glucose tolerance tests and significantly lower basal blood glucose levels (Figure 1d, Supplementary Figure 1c). The KD-10%P group showed comparable basal glucose levels to the chow-fed group (Supplementary Figure 1c) and as above had impaired glucose tolerance (Figure 1d). However, both KD groups were in a ketogenic state with significantly higher blood ketone levels in each (Supplementary Figure 1d). Altering saturated fat content did not modify these effects (Supplementary Figure 2a-d), and varying protein content in carbohydrate-containing chow diets also failed to elicit comparable changes in body weight or glucose metabolism^26^ (Supplementary Figure 2e–g). These data suggest that the effect of a ketogenic diet in B6J mice is a result of the low protein levels and is not secondary to differences in lipid composition or elevated plasma ketone levels.

To assess whether the weight reducing effects of the diet are also evident in obese animals, we placed diet-induced obese (DIO) and leptin-deficient (OB) mice on the KD-5%P and KD-10%P diets. Consistent with data from chow fed wildtype animals, the KD-5%P diet significantly reduced body weight and food intake in DIO and OB mice relative to the KD-10%P diet. (Figure 1e-h, Supplementary Figure 1e-h) (For OB mice, the average final body weight for KD-10%P is 51.78±2.30g, KD-5%P is 35.70±11.95, and chow is 41.88±4.04g. With the 2-way ANOVA test, the adjusted p value for KD-10%P vs. KD-5%P is 0.012, for KD-10%P vs. chow is 0.0944, for KD-5%P vs. chow is 0.3896. And for DIO mice, the average final body weight for KD-10%P is 58.58±8.04g, KD-5%P is 31.51±5.84, and HFD is 52.76±3.89g. With the 2-way ANOVA test, the adjusted p value for KD-10%P vs. KD-5%P is <0.0001, for KD-10%P vs. HFD is 0.4479, for KD-5%P vs. HFD is<0.0001). Moreover, the KD-10%P diet further increased food intake, body weight, and adiposity in OB mice compared to chow fed littermates while body weight was unchanged in DIO mice switched to the KD-10%P (Figure 1e, g). While both diets improved glucose tolerance in DIO and OB mice vs. chow fed controls, the KD-5%P elicited significantly greater improvement in glucose tolerance than the KD-10%P in DIO and OB males with a similar trend in lean females (Figure 1f, h; Supplementary Figure 2h-k).

We next evaluated whether the effects are a result of reduced food intake by pair-feeding animals the KD-10%P group in amounts that matched the (reduced) food intake evident in B6J mice fed the KD-5%P group (Figure 1i). This resulted in a comparable reduction of weight and adiposity suggesting that reduced food intake accounts for the weight loss (Figure 1i). In contrast, the pair-fed KD-10%P group continued to exhibit impaired glucose tolerance with increased fasting glucose levels as well as lower ketone levels (Figure 1j, Supplementary Figure 1i, j). The respiratory exchange ratio (RER) was reduced in both KD groups indicating increased metabolism of fatty acids (Supplementary Figure 3a, b). In addition, the KD-5%P group had significantly lower energy expenditure relative to the KD-10%P group which were similar to chow-fed controls (Supplementary Figure 3c-e). Finally, animals fed a KD-5%P diet showed decreased locomotor and ambulatory activity (Supplementary Figure 3f, g).

In aggregate, these results indicate that the amount of dietary protein is a key determinant of whether lean and obese B6J mice lose weight on a ketogenic diet. Overall, there were two notable differences between the effects of the KD-5%P and KD-10%P in B6J mice: (i). markedly reduced food intake and body weight and (ii). improved glucose tolerance. We next explored the mechanism responsible for each.

### KD-5%P Improves Glucose Tolerance by Limiting Substrates for Gluconeogenesis and Glycogen Synthesis

We first evaluated the basis for the improved glucose metabolism in mice fed the KD-5%P diet by comparing its effects to those of a ketogenic diet (KD) with 5% protein, 5% carbohydrate, and 90% fat (KD-5%P-5%C). This diet resulted in similar effects on body weight, food intake and body composition as the KD-10%P diet, with a similar impairment of glucose tolerance but with significantly lower ketone levels^27^(Figure 2a, b, Supplementary Figure 4a, b). While glycerol from triglycerides can serve as a gluconeogenic substrate, its molar contribution is substantially lower than that of amino acids while fatty acids are even less efficient for producing glucose^28^. This suggested that the difference in glucose production between diets with 5% vs 10% protein might be a result of decreased levels of gluconeogenic precursors in the former. To evaluate a potential difference in the state of activation of metabolic pathways between the groups, we employed COMPASS, an algorithm for flux-balanced analysis mediated inference of metabolic state from RNAseq data^29^. COMPASS revealed that pathways associated with glycolysis/gluconeogenesis and starch and sucrose metabolism had significantly elevated predicted utilization in KD-10%P-fed mice but suppressed in the KD-5%P group, indicating that the substrate levels might be a key driver of the differential response (Figure 2c; Supplementary Figure 4d).

**Figure 2.**
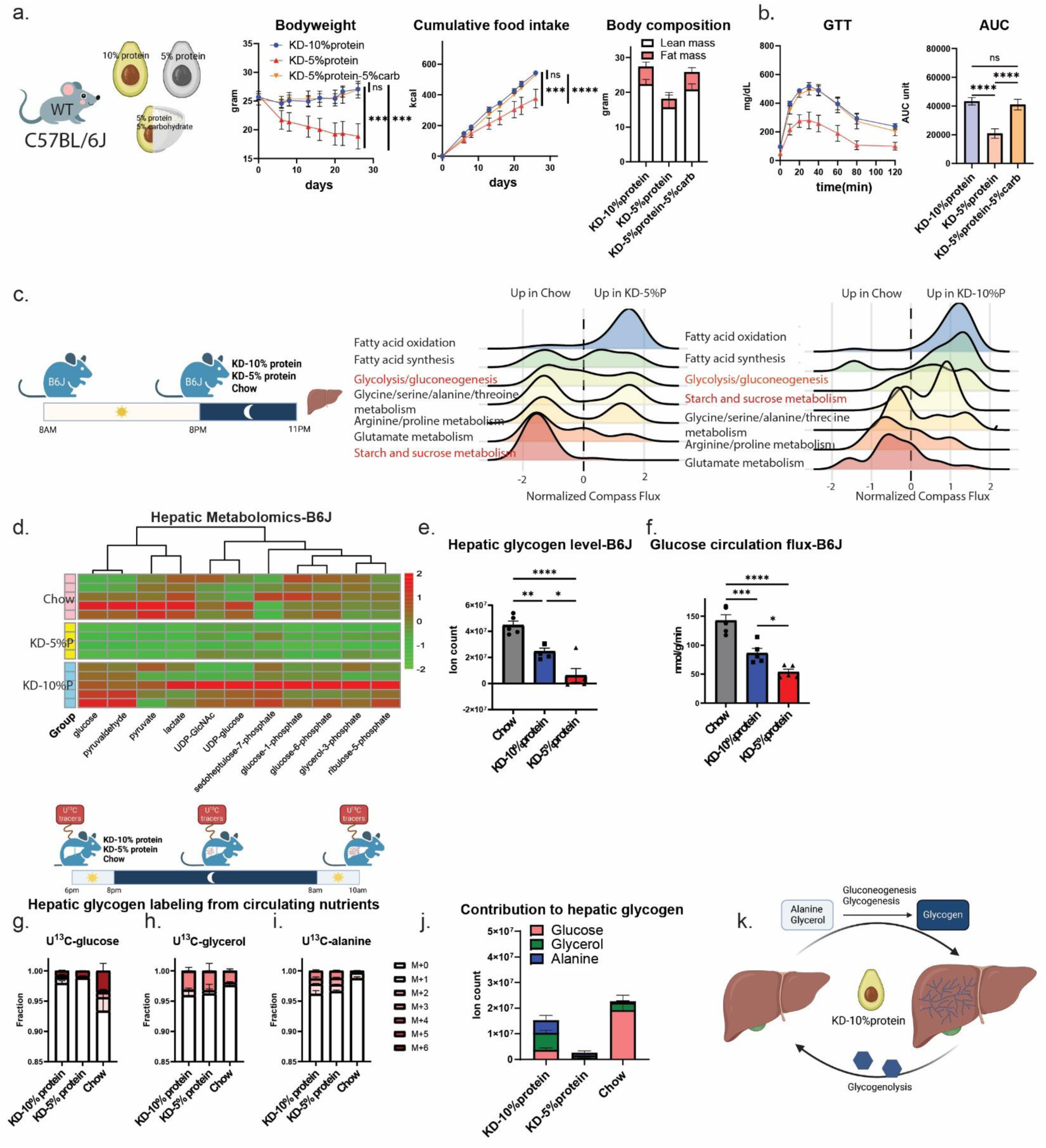
The Improved Glucose Tolerance in the KD-5%P Diet is a Result of Decreased Substrates for Gluconeogenesis and Glycogen Synthesis. a. The schematic of mouse strain, diets and the resulting body weight, cumulative calories consumed, and body composition of WT B6J mice fed with KD-5%P, KD-10%P and KD-5%P-5%C after a 4-week diet challenge. b. Glucose tolerance test (2mg/gram body weight) and calculated area under the curve of WT B6J mice fed with KD-5%P, KD-10%P and KD-5%P-5%C for 4 weeks. c. The schematic of refeeding and harvest livers for transcriptome analysis and resulting compass analysis of hepatic transcriptomes of KD-5%P-fed mice versus chow-fed mice. d. The schematic of LC-MS metabolomics analysis of the mice liver after diets challenge and the heatmap of hepatic glucose metabolites. e. Hepatic glycogen levels of WT B6J mice fed with KD-5%P, KD-10%P and chow. f. Glucose turnover rate measured with in vivo stable isotope tracing in WT B6J mice under KD-5%P, KD-10%P and chow. g. The schematic of glycogen tracing strategy and the resulting glycogen labeling from U^13^C-glucose infusion under KD-5%P, KD-10%P and chow. h. As in g, for U^13^C-glycerol infusion. i. As in g, U^13^C-alanine infusion. j. Calculated contribution from circulating glucose, glycerol and alanine to glycogen under KD-5%P, KD-10%P and chow. k. Proposed mechanism of glucose homeostasis under carbohydrate deprivation. (For a-c, e, f, n = 5 mice for each group; for d, n=5 in KD-10%P and chow fed group and n=4 for KD-5%P group. For alanine tracing, n=3 for KD-5%P, n=3 for KD-10%P, n=5 for chow; for glycerol tracing n=4 for KD-5%P, n=4 for KD-10%P, n=4 for chow; for glucose tracing n=2 for KD-5%P, n=4 for KD-10%P, n=3 for chow. Data are presented as mean ± s.e.m. NS, not significant; *P < 0.05; **P < 0.01; ***P < 0.001; ****P < 0.0001.)

Metabolomic profiling of liver tissue revealed that the KD-5%P group had significantly reduced levels of several substrates for glucose metabolism, whereas KD-10%P-fed mice had metabolite levels comparable to those of chow-fed controls (Figure 2d). Indeed, despite their minimal carbohydrate intake, the KD-10%P-fed mice accumulated relatively normal levels of hepatic glycogen while the KD-5%P-group had a markedly decreased liver glycogen level (Figure 2e, Supplementary Figure 4c). Consistent with prior reports, the mice fed a KD-5%P diet also developed a fatty liver^30^ (Supplementary Figure 4e). Insulin levels were 2-fold lower in the 5%P-fed group relative to the KD-10%P-fed which were equivalent to chow fed mice and glucagon levels were similar among the 3 groups (Supplementary Figure 5a, b)^31,32^. However, even in the absence of insulin following streptozotocin (STZ) treatment, B6J mice fed the KD-10%P diet maintained higher body weight compared to those fed the KD-5%P diet (Supplementary Figure 5c-i). This is in contrast to a prior report^17^ suggesting that insulin does not account for the differential effects of the two KDs.

To further evaluate the possible role of the lower level of gluconeogenic metabolites in the KD-5%P group, we next measured glucose turnover and the incorporation of different substrates into glycogen. We measured glucose turnover after 3 weeks on the two diets by infusing U¹³C-glucose. Mice fed the KD-10%P exhibited significantly higher circulating glucose flux than the KD-5%P group, suggesting an increased rate of glucose utilization and production with higher levels of hepatic glycogen stores^33^ (Figure 2f, Supplementary Figure 6a). To further investigate this, we then measured the incorporation of three ¹³C-labeled substrates-glucose, glycerol, and alanine- into glycogen in B6J mice fed the two diets. Each substrate was intravenously infused over 16 hours following the onset of the light cycle and labeling of glycogen was assessed using mass spectrometry. In chow-fed mice, glucose accounted for over 40% of glycogen synthesis, whereas glycerol and alanine together contributed about 10% (Figure 2g-i, Supplementary Figures 6b, d, g). Consistent with the results showing reduced glucose turnover, less than 15% of the infused glucose was used to synthesize glycogen on either of KDs while glycerol contributed ∼26% in mice fed KD-10%P and 16% in mice fed the KD-5%P (Figure 2g-i, Supplementary Figures 6d-f). Finally, alanine was incorporated into glycogen and glucose to a similar extent as glycerol (20% in KD-10%P and 16% in KD-5%P) (Figure 2h, i; Supplementary Figures 6g-i). After normalizing for the level of infusate labeling (the percentage labeling of tracer in serum), the amount of alanine incorporation into glycogen was similar to that of glycerol (Figure 2j). Alanine represents only 4.4% of the total protein (Supplementary Figure 6j) suggesting that the total amount of protein incorporated into glycogen greatly exceeds that of glycerol. Furthermore, the KD-5%P-fed mice consume only ∼30% as much protein (50% less protein in diet plus a 40% reduction in food intake) as the KD-10%P-fed group further contributed to their lower glucose and glycogen levels and underscoring the critical contribution of dietary protein for maintaining normal glucose levels in these studies.

In summary, substituting carbohydrates for protein in the ketogenic diet, together with COMPASS analysis of hepatic RNAs, metabolite profiling, and flux studies, indicate that reduced gluconeogenic/glycogenic substrates underlie the lower rate of glucose metabolism in KD-5%P mice, whereas increased dietary protein intake in KD-10%P-fed mice sustains glucose production through amino acid–derived carbon flux (Figure 2k). We next turned our attention to the mechanisms responsible for the reduced food intake seen in animals fed the KD-5%P diet.

### GDF15 Partially Accounts for the Reduced Food Intake of Mice Fed the KD-5%P Diet

We next measured the RNA levels for liver genes encoding secreted peptides using RNAseq and consistent with prior reports found increased levels of several anorexigenic peptides in B6J mice fed the KD-5%P relative to KD-10%P. These included GDF15^19^ and FGF21^25^ as well as, CCL2^34^, LCN2^35^, and Apoa4^36^ (Figure 3a). Serum ELISA assays confirmed significantly increased circulating levels of FGF21 and GDF15 in B6J mice fed the KD-5%P while the levels were unchanged in the KD-10%P group (Average GDF15 level for KD-10%P is 657.59±114.18 ng/ml, KD-5%P is 1459.886±313.0592 ng/ml and for chow is 436.5507± 207.4694 ng/ml. With the ordinary one-way ANOVA test, the adjusted p value for KD-10%P vs. KD-5%P is 0.0497, for KD-10%P vs. chow is 0.7809, for KD-5%P vs. chow is 0.0142.) (Figure 3b, c). FGF21 has been linked to an increased energy expenditure and water intake in animals fed ketogenic diets^9,25,37^. However, FGF21-KO mice fed the KD-5%P still exhibited reduced food intake, body weight, and improved glucose tolerance than KD-10%P fed group (Supplementary Figures 7a-e), indicating that the differential effects of the KD-5%P are independent of FGF21.

**Figure 3.**
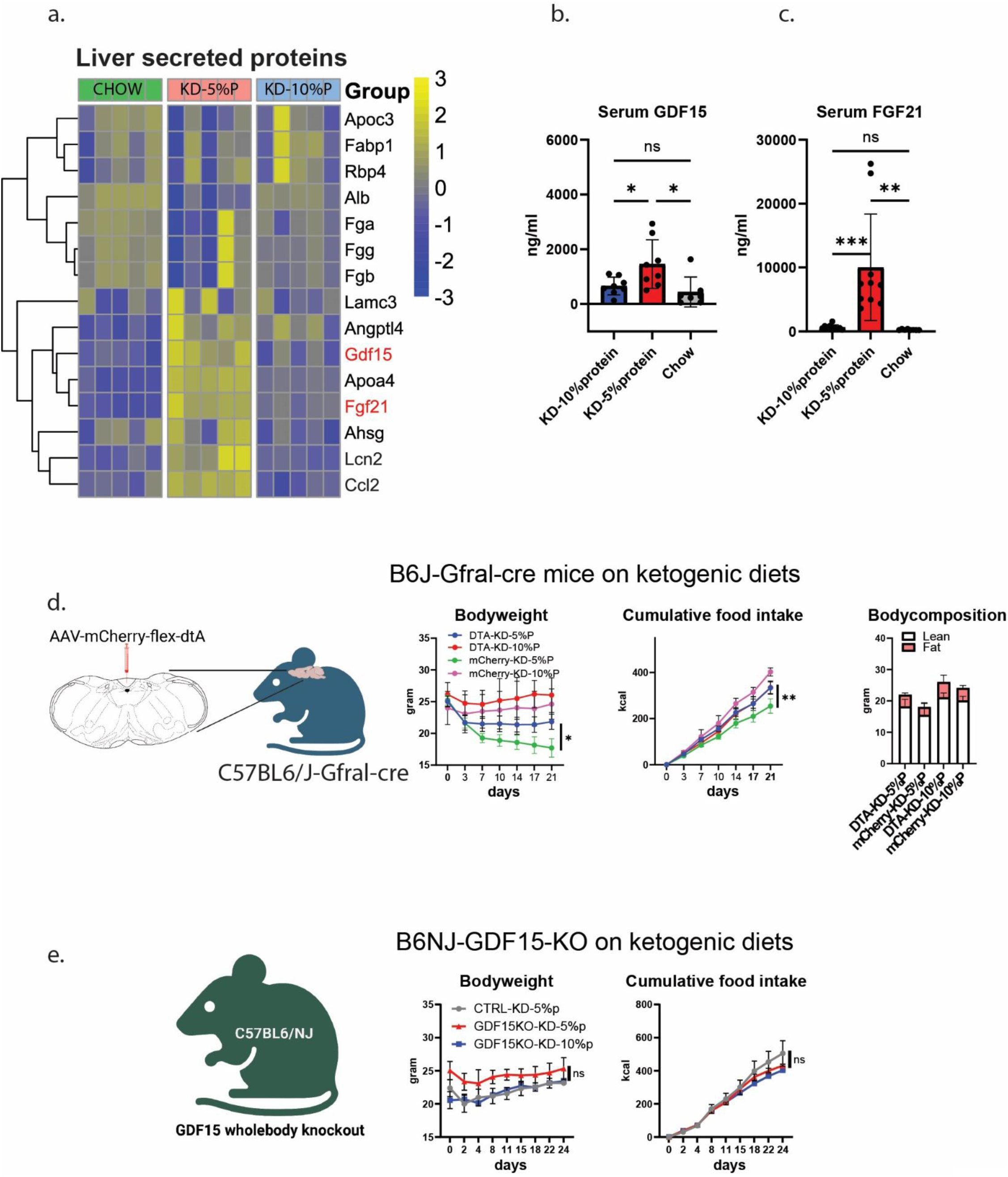
GDF15 Partially Accounts for the Reduced Food Intake of Mice Fed the KD-5%P Diet. a. Heatmap of secreted protein expression in liver on KD-5%P, KD-10%P and chow. b. Circulating GDF15 levels of WT B6J mice fed with KD-5%P, KD-10%P and chow. c. Circulating FGF21 levels of WT B6J mice fed with KD-5%P, KD-10%P and chow. d. The schematic of ablating GFRAL-neurons in Area Postrema and the resulting body weight, cumulative calories consumed (kcal), and body composition of WT B6J mice under KD-5%P or KD-10%P. e. Body weight and cumulative calories consumed of B6NJ-GDF15 whole-body knockout and control mice under KD-5%P or KD-10%P. (n = 3 mice for DTA fed with KD-10%P and mCherry with KD-10%P; n=7 for DTA fed with KD-5%P and n=9 for mCherry fed with KD-5%P. GDF15-KO experiment has n=2 in each group. Data are presented as mean ± s.e.m. NS, not significant; *P < 0.05; **P < 0.01; ***P < 0.001; ****P < 0.0001.)

We then investigated the possible contribution of GDF15 by first ablating neurons expressing its receptor, GFRAL, in the Area Postrema (AP) and Nucleus of the Solitary Tract (NTS) of the brain stem, the principal site of GDF15 action^19,38,39^. We injected an AAV expressing a cre-dependent DTA^40^ into the AP/NTS of GFRAL-cre mice, and found that mice with an ablation of GFRAL neurons showed an attenuated response to a KD-5%P with increased food intake, body weight, and lean mass and impaired glucose tolerance compared to control mice fed this diet.(Average final body weight for DTA fed with KD-10%P is 26.03±2.75g, KD-5%P is 21.84±1.19g. For mCherry fed with KD-10%P is 24.62±1.83g, KD-5%P is 18.14±1.34. With the 2-way ANOVA test, the adjusted p value for DTA-KD-5%P vs. mCherry-KD-5%P is 0.0454, DTA-KD-5%P vs. DTA-KD-10%P is 0.1838, DTA-KD-5%P vs. mCherry-KD-10%P is 0.5364, mCherry-KD-5%P vs. DTA-KD-10%P is 0.0009, mCherry-KD-5%P vs. mCherry-KD-10%P is 0.0085. DTA-KD-10%P vs. mCherry-KD-10%P is 0.9446.) (Figure 3d, Supplementary Figures 8a). These findings indicate that GDF15–GFRAL signaling contributes to, but does not fully account for, the metabolic effects of KD-5%P.

To further confirm the contribution of GDF15, we evaluated the response of GDF15-KO mice to the KDs. Surprisingly, in this case we found that both the knockout mice and wild type littermate controls showed similar responses to the KD-5%P and KD-10%P diets with neither group showing a reduced food intake, body weight or altered glucose levels (Figure 3e, Supplementary Figure 8b). Upon closer inspection, we realized that the GDF15-KO line is maintained on the C57BL/6NJ (B6NJ) background rather than the B6J strain used in our initial studies. We then compared the metabolic response of a different cohort of age-matched B6J and B6NJ male mice to the two ketogenic diets. While B6J mice showed the same response to the KD-5%P diet as above, the weight of B6NJ mice was similar in groups fed the KD-5% and KD-10%P diets with similar food intake (Figure 4a). As before, while the GDF15 levels increased in B6J mice, it failed to increase in the B6NJ mice fed with KD-5% (Figure 4b). The B6NJ mice fed with the KD-5%P and KD-10%P diets also had normal plasma glucose with higher ketones and did not develop a fatty liver in contrast to the B6J mice on the KD-5%P (Supplementary Figures 8c-g). These strain-specific differences were consistent across sexes (Supplementary Figures 8h-l). To further assess strain dependency, we examined DBA/2J and FVB/NJ mice and found that, similar to B6NJ, neither strain showed differential responses to KD-5%P versus KD-10%P (Supplementary Figures 9a-f).

**Figure 4.**
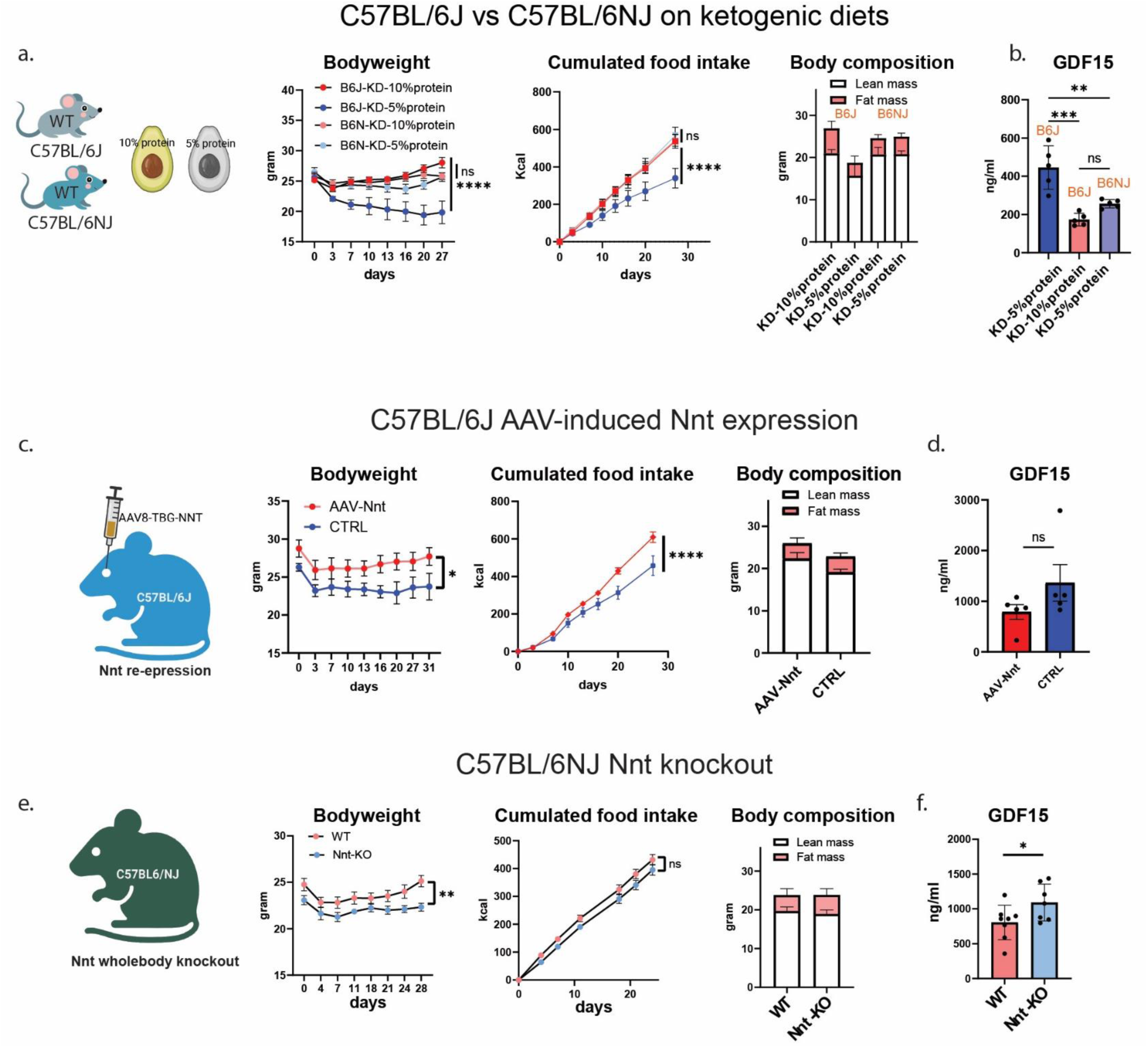
Nnt-Gdf15 Partially Mediates the Strain-specific Differences in the Response to the KD-5%P Diet. a. Body weight, cumulative calories consumed, and body composition of WT B6J and B6NJ mice under KD-5%P or KD-10%P. b. Serum GDF15 levels of WT B6J and B6NJ mice under KD-5%P or KD-10%P. c. The schematic of hepatic AAV-induced Nnt expression in B6J mice and the resulting body weight, cumulative calories consumed, and body composition of both groups under KD-5%P. d. Serum GDF15 levels of AAV-induced Nnt expression and control mice under KD-5%P. e, f. As in c, d, for B6NJ Nnt whole-body knockout mice. (n = 5 for each B6J and B6NJ group, n=5 for AAV-induced Nnt expression and control group. n=8 for Nnt-WT, n=7 for Nnt-KO. Data are presented as mean ± s.e.m. NS, not significant; *P < 0.05; **P < 0.01; ***P < 0.001; ****P < 0.0001.)

The B6J and B6NJ are two sub-strains that diverged in 1950 with the B6J strain being maintained at Jackson(J) and the B6NJ mice maintained at NIH(N). Whole genome sequencing has revealed that ∼.1% of the genome has diverged between the two strains^21^. Among these small numbers of changes, B6J mice carry a mutation in the Nnt gene while B6NJ, DBA and FVB/NJ mice are wild types at this locus. Nicotinamide nucleotide transhydrogenase (Nnt) is a mitochondrial enzyme required for generating NADPH from NADH in mitochondria^41^, and a differential response to ketogenic diets in these two strains was previously attributed to differences insulin-secretion^42^. In light of our findings in streptozotocin treated mice (see Supplementary Figure 5 above) that the effects of the diets were independent of insulin we considered the possibility that a mutation in Nnt led to weight loss in B6J mice fed the KD-5%P. B6J also carries a mutation in the Nlrp12 gene^21,43^, which also contributed to the different responses of the two strains, see below.

### NNT-GDF15 Axis Partially Mediates Strain-specific Differences in the Response to the KD-5%P Diet

Nnt plays a pivotal role in blunting the response to mitochondrial and oxidative stress^21,41^. Because GDF15 expression is induced by stress^19,38^, we hypothesized the anorectic response of B6J mice to the KD-5%P diet was a result of activation of stress pathways. To confirm the role for Nnt in this response, we injected an AAV expressing Nnt retro-orbitally which leads to re-expression in liver and elsewhere. B6J mice fed the KD-5%P that received the Nnt AAV showed a significantly increased food intake, body weight, normal fasting glucose levels with a further impairment of glucose tolerance and a decreased GDF15 level relative to B6J mice receiving a control virus (Figure 4c, d, Supplementary Figures 9g, h) Similarly, B6NJ mice with an Nnt knockout now showed a similar response as B6J mice fed the KD-5%P diet with reductions in food intake, body weight, blood glucose, impaired glucose tolerance and increased GDF15 level (Figure 4e, f, Supplementary Figure 9i). However, the B6NJ Nnt KO mice showed smaller differences than B6J mice on the two KDs raising the possibility that additional factors between the two strains may also contribute. Nonetheless, these findings suggest that the Nnt mutation contributes significantly to the effect of a low protein KD-5%P diet to reduce food intake and body weight via GDF15.

### NLRP12-LCN2 Contributes to the Strain-specific Differences in the Response to the KD-5%P Diet Independent of GDF15

The comparative transcriptomic analyses also revealed that the KD-5%P also induced liver genes enriched for immune pathways in C57BL/6J (B6J) mice, including increased expression of the Lcn2, Ccl2, genes as well as those for other inflammatory mediators (Figure 3a). These responses were not evident in C57BL/6NJ (B6NJ) mice (Figure 5a, b). Several of these factors are known to be regulated by Nlrp12, an anti-inflammatory regulator^43^, which, as mentioned, is also mutant in B6J mice^21,43^. Plasma levels of LCN2 and CCL2 were also significantly elevated in B6J mice fed the KD-5%P but not in B6NJ mice (Figure 5c). Similar to the effect of GDF15, a knockout of Lcn2 in B6J mice partially rescued the hypophagia and weight loss seen in B6 animals fed the KD-5%P (Figure 5d, Supplementary Figure 10c), suggesting that LCN2 also contributes to the anorexic effect of KD-5%protein in B6J mice. While serum CCL2 was also selectively increased in B6J mice fed with KD-5%P, a Ccl2 knockout failed to alter food intake or body weight (Supplementary Figure 10a-c).

**Figure 5.**
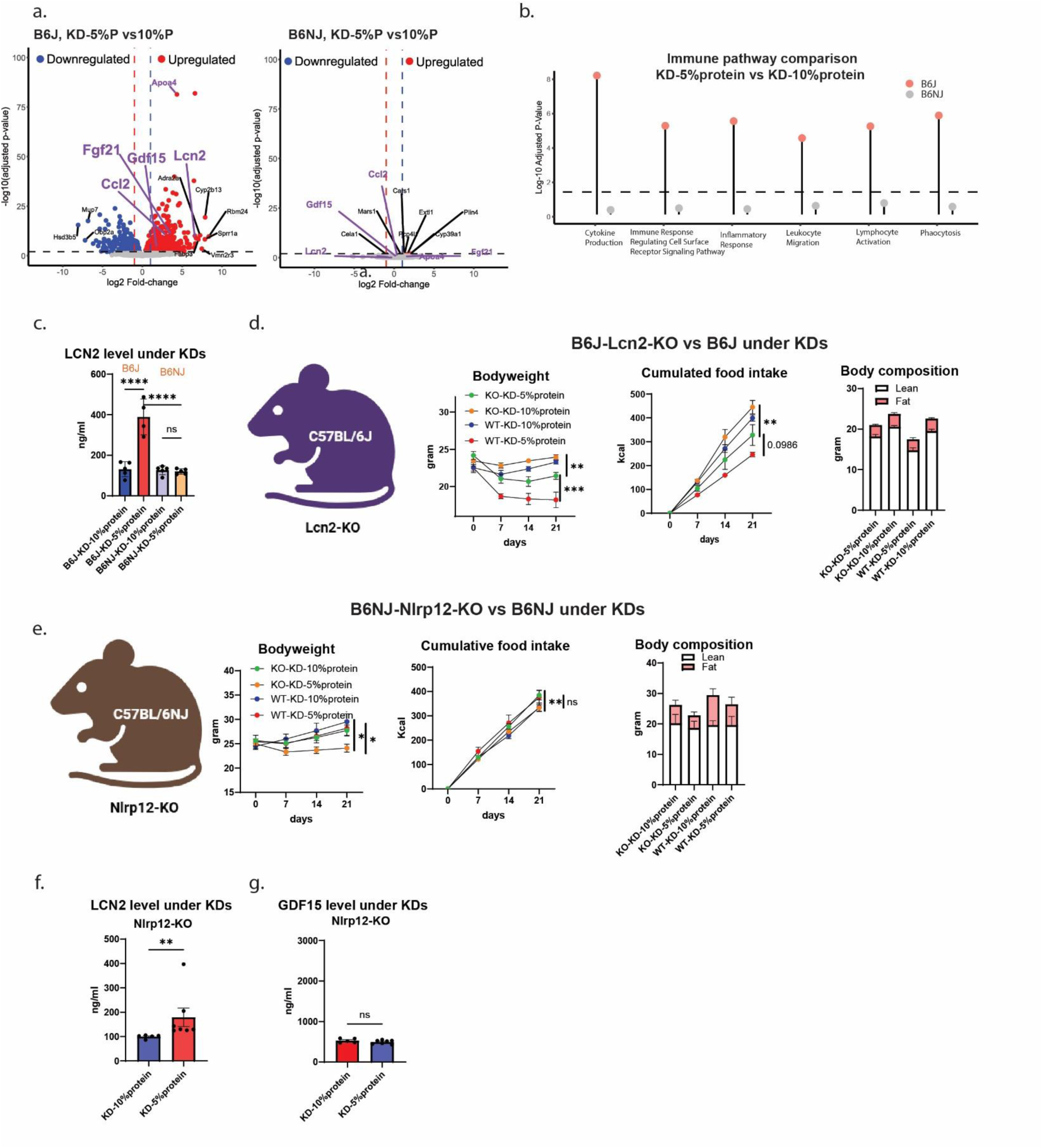
Nlrp12-Lcn2 Contributes to the Strain-specific Differences in the Response to the KD-5%P Diet Independent of GDF15. a. Volcano plot of hepatic mRNA expression of B6J and B6NJ mice fed with KD-5%P versus KD-10%P at ad lib fasted state. b. Immune response pathway analysis between KD-5%P versus KD-10%P in B6J and B6NJ mice. c. Serum LCN2 levels of WT B6J and B6NJ mice under KD-5%P or KD-10%P. d. Body weight, cumulative calories consumed, and body composition of B6NJ-Nlrp12 whole-body knockout and littermate mice under KD-5%P or KD-10%P. e. Body weight, cumulative calories consumed (kcal) and body composition of B6J-Lcn2 whole-body knockout and littermate mice under KD-5%P or KD-10%P. f. Serum LCN2 levels of B6NJ-Nlrp12 whole-body knockout mice under KD-5%P or KD-10%P. g. Serum GDF15 levels of B6NJ-Nlrp12 whole-body knockout mice under KD-5%P or KD-10%P. (For a, b and d, n = 5 for each B6J and B6NJ group; n=10 for Nlrp12-KO under KD-5%P, n=8 for Nlrp12-KO under KD-10%P, and n=5 for both WT litter mate under 2 diets. For B6J-Lcn2 KO experiment, n=7 for KO under KD-5%P, n=5 for KO under KD-10%P and n=4 for WT under both diets. For serum Lcn2 and GDF15 measurement in B6NJ-Nlrp12-KO mice, n=5 for KD-10%P group and n=7 for KD-5%P group. Data are presented as mean ± s.e.m. NS, not significant; *P < 0.05; **P < 0.01; ***P < 0.001; ****P < 0.0001.)

Because LCN2 is induced by inflammation^44^ and NLRP12 is an anti-inflammatory regulator^45^, we studied B6NJ mice with an Nlrp12 knockout. B6NJ mice with Nlrp12 mutations now lost significant amounts of body weight with decreased adiposity and food intake—phenotypes not observed in wild-type B6NJ mice (Figure 5e), showing that the NLRP12-LCN2 axis contributes to the anorexia effect induced by KD-5%P. We also found that the KD-5%P diet increased circulating LCN2 but not GDF15 levels in Nlrp12-deficient B6NJ mice (Figure 5f, g), indicating that loss of Nlrp12 selectively regulates anorexigenic signals independently of the GDF15 stress pathway.

Together, these results identify an NLRP12-LCN2 immune pathway that also contributes to strain-specific sensitivity to KD-5%P and operates in parallel with stress-associated mechanisms that activate GDF15, providing a mechanistic explanation for the susceptibility of B6J mice to effects of very low-protein ketogenic diets.

### Oxidative Stress Integrates Stress and Immune Pathways to Drive the Response of B6J Mice to the KD-5%P Diet

Having identified distinct stress-associated (NNT–GDF15) and immune-associated (NLRP12–LCN2) pathways that are associated with the sensitivity of B6J mice to the KD-5%P, we tested the role of each using a pharmacologic approach. Pathway analysis of the differentially expressed genes identified markers of both an integrated stress response (ISR) and an oxidative stress response solely in the livers of B6J fed the KD-5%P (Supplementary Figures 11a-c). Evidence for activation of the ISR included induction of Atf4 mRNA and many of its target genes including Ddit3, Lonp1, Hspa9 and Gdf15 (Figure 6a). Additionally, we observed elevated levels of Nrf2, the master regulator of antioxidant responses, along with its target genes Gpx3, Hmox1, Sod3, and Nqo1, which mediate oxidative stress defense (Figure 6b). These findings suggested that KD-5%P engages in overlapping stress programs downstream of impaired redox metabolism in B6J mice.

**Figure 6.**
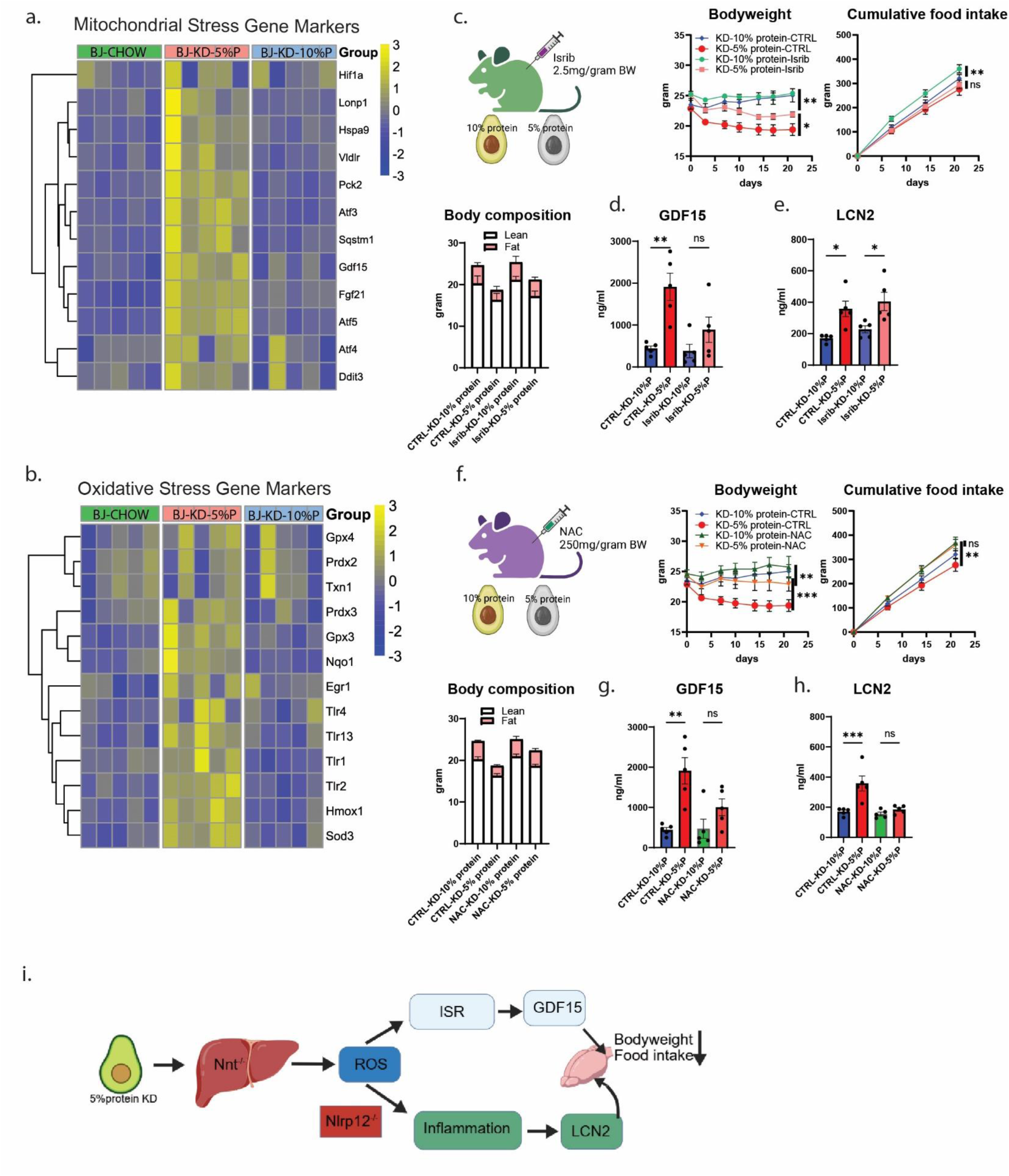
Oxidative Stress Induced Hepatic Inflammation Accounts for the Response of B6J Mice Fed the KD-5%P. a. Heatmap of mitochondrial stress gene markers of B6J mice fed with KD-5%P, KD-10%P and chow. b. Heatmap of oxidative stress gene markers of B6J mice fed with KD-5%P, KD-10%P and chow. c. The schematic of Isrib i.p. injection in B6J mice and the result body weight, cumulative food intake (Kcal) and body composition of both groups under KD-5%P or KD-10%P. d. Serum GDF15 levels of B6J mice treated with Isrib or Vehicle under KD-5%P or KD-10%P. e. Serum LCN2 levels of B6J mice treated with Isrib or Vehicle under KD-5%P or KD-10%P. f-h. As in c-e, for NAC treatment, control group is the same data. i. The schematic of the mechanism on how KD induces body weight loss (n = 5 mice for each group. Data are presented as mean ± s.e.m. NS, not significant; *P < 0.05; **P < 0.01; ***P < 0.001; ****P < 0.0001.)

We tested the contribution of ISR by administering Isrib, an ISR inhibitor, which stabilizes the EIF2 complex and reduces the expression of ATF4 regulated genes^46^(Figure 6c). B6J mice fed the KD-5%P diet and treated with Isrib (2.5 mg/kg, i.p., every other day) showed a small but significant increase in body weight compared to vehicle treated controls with no significant changes in food intake or glucose tolerance (Average body weight of Isrib group on KD-10%P is 25.37±0.36g and on KD-5%P is 21.88±0.42g; of control group on KD-10%P is 24.66±1.10g and on KD-5%P is 19.28±0.88g. With the 2-way ANOVA test, the adjusted p value for KD-5% protein-Isrib vs. KD-5% protein-CTRL is 0.0349.) (Figure 6c; Supplementary Figure11 e-g). Isrib administration also led to a small reduction in circulating GDF15 levels but not LCN2 (Figure 6d, e). These results suggest that the activation of the ISR may contribute to the induction of GDF15 in B6J mice fed the KD-5%P but that this effect was insufficient to fully account for the phenotype.

We next evaluated the possible contribution of oxidative stress by treating B6J mice fed KD-5%P with the antioxidant N-acetylcysteine (NAC) a potent antioxidant and anti-inflammatory agent as well^47–50^. Treatment of B6J mice fed the KD-5%P diet with NAC (i.p. 250 mg/kg BW daily) significantly increased body weight, food intake, decreased glucose tolerance, and reduced GDF15 levels to levels similar to those seen in B6J mice fed the KD-10%P diet and B6NJ mice fed the KD-5%P (Figure 6f, Supplementary Figures 11e-g). Moreover, in contrast to ISR inhibition, NAC simultaneously suppressed both GDF15 and LCN2 levels (Figure 6g, h), suggesting that it attenuates inflammatory signaling downstream of oxidative stress. Combined administration of ISRIB and NAC did not further improve metabolic outcomes beyond NAC alone (Supplementary Figure 11i–l), indicating that oxidative stress is likely to be upstream of both ISR activation and immune amplification.

By simultaneously suppressing GDF15-mediated stress signaling and LCN2-associated inflammatory anorexia, antioxidant treatment effectively rescued the phenotypes, explaining its superior efficacy relative to ISR inhibition alone. Together, these findings identify oxidative stress as a central integrator of the anorectic response of C57BL/6J mice to a KD-5%P-induced stress response.

## Discussion

In this study, we found that protein content is a primary determinant of the ability of a ketogenic diet (KD) to reduce weight in B6J mice. Thus, in C57BL/6J (B6J) mice, a low-protein ketogenic diet (KD-5%P) induces profound hypophagia, weight loss, and hypoglycemia, whereas the same carbohydrate - free diet containing higher protein content promotes weight gain. Surprisingly, we found that the effect of this same diet is not evident in C57BL/6NJ (B6NJ) mice which only carry a small number of differences relative to B6J mice. This finding demonstrated that genetic background strongly modulates the response to the low protein ketogenic diet that we employed.

Our data show that this divergence arises from coordinated activation of stress and immune pathways. B6J mice harbor loss-of-function mutations in Nnt, which regulates mitochondrial redox balance, and Nlrp12, an anti-inflammatory regulator^21,41,43,51^. In the setting of a low-protein KD, impaired NNT function leads to activation of hepatic oxidative stress and the integrated stress response, culminating in induction of the anorexigenic factor GDF15. Pharmacologic suppression of oxidative stress normalized food intake, body weight, and circulating GDF15 levels, establishing oxidative stress as a key upstream driver of KD-5%P–induced anorexia. This effect is also evident in DIO and OB mice showing that activation of this pathway can override the hyperphagia that develops in these obese animals, both of which have defects in leptin signaling.

While this stress-associated signaling pathway contributes to weight loss in B6 mice fed the KD-5%P, our data further reveal that immune dysregulation amplifies the anorectic response. Loss of Nlrp12 results in exaggerated hepatic inflammatory responses and selective induction of the immune-derived anorexigenic factor LCN2^22,35^. Genetic deletion of Nlrp12 rendered otherwise resistant B6NJ mice sensitive to KD-5%P–induced weight loss without increasing GDF15. These findings indicate that an NLRP12–LCN2 immune axis that operates in parallel with the NNT–GDF15 stress pathway, explaining why alterations of either pathway alone only have a partial effect while reducing the activity of both pathways with antioxidant treatment suppresses both oxidative stress and inflammation and is substantially more effective.

The mechanism by which a ketogenic diet induces weight loss has been intensively studied and helps to reconcile conflicting reports regarding the mechanisms. While the PPARα-FGF21 axis was first identified because it is activated by a KD, our results indicate that it plays little or no role to induce a state of negative energy balance^9,25,52^. Recently, a novel ketone metabolite β-hydroxybutyryl-phenylalanine (Ket-Phe) was identified and shown to have weight-reducing effects, though only at supraphysiologic concentrations^23^. While we find increased levels of ketone bodies in both B6J and B6NJ mice fed the KD-5%P, the fold change is relatively small, suggesting that it too is unlikely to account for the negative energy balance that we observe.

Instead, and consistent with prior reports, we find that GDF15 contributes to a state of negative energy balance in mice fed a KD^19,38^. GDF15 reduces food intake as well as inducing nausea in several clinical settings including cancer cachexia^53^, inflammation, and pregnancy^54^. Our data also links its activation to a mutation in NNT and activation of an oxidative stress response. While GDF15-GFRAL signaling contributes significantly to KD-induced anorexia, we found that neither GFRAL neuron ablation nor a GDF15 knockout normalized food intake suggesting that additional mechanisms contribute. Our identification of LCN2 as a stress-independent anorexigenic factor provides a mechanistic explanation for this residual effect of the diet and indicates that immune-derived signals cooperate with stress pathways in a diet and strain dependent manner to regulate energy balance.

It is not clear why the diet activates these oxidative and integrative stress pathways and an immune response, but we hypothesize that it is a result of the markedly increased level of fatty acid oxidation in the absence of carbohydrate and protein. Fatty acid oxidation is known to generate ROS^55,56^ which in addition to causing a fatty liver^41,57^ can activate stress and inflammatory responses in liver^58^. In animals fed with KD-5%P, the principal source of energy is fatty acids with reduced glucose utilization as evidenced by the low RER and the decreased glucose turnover. In contrast to our data, a previous study reported that an Nnt mutation only had a modest effect on the response of B6J mice to a high fat diet which was attributed to differences in insulin secretion^42^. In contrast, in studies of mice treated with streptozotocin, we find that the response to ketogenic diets lacking carbohydrates is independent of insulin secretion. We confirmed that oxidative stress is a major contributor to KD-5%P sensitivity by treating B6J mice with N-acetylcysteine (NAC), a well-established antioxidant and anti-inflammatory compound. NAC essentially normalized food intake, body weight, and circulating GDF15 and LCN2 levels. In contrast, although KD-5%P also induced the integrated stress response (ISR), inhibition of this pathway produced only partial rescue, and combined treatment with NAC and ISRIB was no more effective than NAC alone. Because oxidative stress can activate both ISR signaling and inflammatory pathways, the lack of synergy suggests that oxidative stress occupies a more central, upstream position in driving the anorectic and metabolic effects of KD-5%P. Importantly, NAC also suppressed immune-associated outputs, consistent with its ability to attenuate both stress- and inflammation-driven anorexigenic signals^59^.

The two diets also had differential effects on glucose metabolism in B6J mice. The maintenance of glucose homeostasis during prolonged carbohydrate restriction is a profound metabolic challenge^17,33^ and our findings suggest that in the absence of carbohydrate, gluconeogenesis from amino acids, but not solely glycerol, is required to maintain glucose and glycogen levels. Thus, we find that when dietary protein falls below a critical level in the absence of ingested carbohydrate, severe hypoglycemia develops. However, we also find that because B6NJ mice are not hypophagic (compared to B6J), they do not develop hypoglycemia even when fed with the KD-5%P. The data thus indicate that in the absence of carbohydrate there is a critical threshold of total protein intake that is necessary to maintain blood glucose^26^ further suggesting that it is the combination of both decreased protein content and food intake that triggers hypoglycemia in B6J mice fed the KD-5%P. This also raises the possibility that in animals fed the KD-10%P, glucose is primarily diverted for the maintenance of serum glucose level rather than as a source of fuel energy^60^. This is consistent with our finding that there is higher glycogen levels in animals fed the diet with a higher protein level as glycogen can provide as a reservoir to stabilize circulating glucose levels.

The extent to which low-carbohydrate diet or ketogenic diets reduce weight in humans are variable and the subject of ongoing debate^12,15,61^. Our results suggest that interaction between macronutrient composition and genetic variation in stress and inflammatory pathways may similarly influence responses to ketogenic diets in humans. Loss-of-function mutations in Nnt have been identified in patients with familial glucocorticoid deficiency and are associated with impaired mitochondrial redox homeostasis and increased oxidative stress^62^, while variants in Nlrp12 are linked to autoinflammatory syndromes characterized by dysregulated innate immune signaling^63^. Although these conditions are rare, they show defects in oxidative stress buffering and inflammatory restraint can profoundly alter systemic physiology. More broadly, individuals that are sensitive to oxidative stress (G6PD deficiency^64^) or inflammation (other autoimmune diseases) may respond to ketogenic diets more strongly and our findings emphasize the need to design precision nutritional interventions.

Overall, while our findings are consistent with several publications each reporting different metabolic responses of the two B6 sub-strains^42,51,65^, our results provide a comprehensive mechanistic explanation. Thus, our data integrates a number of previous findings by showing that: (i). The weight loss seen in C57BL/6J mice fed a ketogenic diet is a result of decreased protein content in the diet with little or no effect of altering the types of lipids. (ii). This decreased protein content in combination with an Nnt mutation in C57Bl/6J mice results in an increased oxidative stress response in liver rather than altering insulin secretion from β-cells. (iii). The oxidative stress response leads to increased expression of GDF15 in liver and a role for GDF15 was confirmed by ablating GFRAL in brain stem which significantly decrease the anorectic effect of the low protein diet. (iv). Altered regulation of immune factors due to Nlrp12 mutation amplifies anorexia through immune-derived factors such as LCN2. (v). Antioxidant treatment suppresses both stress- and immune-mediated pathways and attenuates the increase of GDF15 and Lcn2 thus explaining its superior efficacy. (vi). The effects of KD-5%P on glucose metabolism are substrate driven and largely independent of β-cell function or insulin production^42,51,65^.

In summary, our findings establish the mechanism in which macronutrient composition and genetic background jointly shape ketogenic diet outcomes through coordinated stress and immune pathways. By identifying stress-immune axis as a convergent driver of anorexia and metabolic remodeling, this work provides a unifying explanation for previously inconsistent observations in ketogenic diet studies in both mice and humans^15,19,23,66^. More broadly, these results underscore the importance of considering both genetic susceptibility and inflammatory state when designing dietary interventions and support the development of precision nutritional strategies for metabolic disease.

## Methods

### Animals

Wild-type C57BL/6J (#000664), C57BL/6NJ (#005304), FVB/NJ(#001800), DBA/2J (# 000671), Fgf21-KO (B6.129Sv(Cg)-Fgf21^tm1.1Djm^/J, #033846), Gfral-cre(B6;SJL-Gfral^em1(cre)Rsy^/J, #036750), *Gdf15^nuGFP-CE^(C57BL/6 Gdf15tm1(cre/ERT2)Amc/J,* 034497) were obtained from Jackson Lab. OB F1 were obtained from Jackson Lab, then bred in lab, from crossing male OB and female ob/+(#000632). Wild-type inbred mice are purchased at 7-wk-old while genetically modified mice are bred in lab. If not specifically indicated, all lean animals were kept at (20°C) a 12 h light-dark cycle (lights on at 7am to 7pm) and on a standard-chow diet (PicoLab Rodent Diet 205053) until 8-wk-old to perform experiments. Diet-induced obese mice are purchased 6-wk-old C57BL/6J mice fed with a high-fat diet in house (HFD, Research Diets, Cat# D12492, Rodent Diet With 60 kcal% Fat) for 16 weeks. Insulin-withdrawal assay was performed on purchased 7-12wk-old C57BL/6J mice. Mice were given one dose of streptozotoxin (200 mg/kg body weight) i.p. and monitored closely for 12 hours providing 10% sucrose water. Replace sucrose water with normal water on day 3 and mice were hyperglycemic until day10, then experiments are performed. Jugular vein catheterization was performed in-house following the protocol^67^. All experiments were performed under approved animal protocols, both male and females were used. All experiments were done in an age and gender-matched way.

### Diets and feeding experiments

Ketogenic diets are customized from Inotive with the following catalog numbers: KD-10%P (#96355), KD-5%P (#210657), KD-10%P-MUFA (#210574), KD-10%P-SAFA (#210575), KD-5%P-SAFA (#220243), KD-5%P-5%C (#230393). High carbohydrate chow diets are customized from TestDiet with the following catalog numbers: Chow-20%P (#5S4D), Chow-10%P (#9G55), Chow-5%P (#5Z6U). All detailed nutritional values are provided in the raw data files.

All mice were single-housed and given a specific diet at the age of 8 weeks if not specifically noted, ad libitum feed for 3 weeks, body weight, food intake, blood glucose and ketone are measured weekly or biweekly. GTT and MRI are performed in the 4^th^ week. The pairfeeding experiment was conducted by measuring the daily food intake of the KD-5%P group which shows lower food intake and manually feeding that same amount of KD-10%P to the other group. 2 control groups are all ad lib fed with either KD-5%P or KD-10%P.

### Glucose, ketone measurement and glucose tolerance test

Blood glucose was measured by tail vein sampling using a Breeze2 glucometer (Bayer SKU: breeze2meter UPC: 301931440010). Blood ketone levels are measured by tail vein sampling using a GK+ Keto-Mojo Meter. All animals were fasted overnight before GTT. Lean animals were given 2 mg/kg body weight glucose in saline, while obese or diabetic animals were given 0.5 mg/kg body weight glucose in saline. For GTT, glucose is measured at 0, 10, 20, 30, 40, 60, 90 and 120 min.

### Body weight and body composition monitoring

Total body mass was measured by a CGOLDENWALL high precision lab scale. Body fat mass was measured by MRI in live mice using an Echo-MRI 100H (EchoMRI, LL). Behaving mice are acclimated before MRI testing and MRI machine are calibrated before use.

### Metabolic cages

Singly-housed wild-type C57BL/6J mice were control or ketogenic diets for 2 weeks before sending into the metabolic cages, which are climate-controlled (temperature: 22 °C; humidity: 55%; 12-hour light–12-hour dark cycle) automated home cage phenotyping system (TSE) with ad libitum access to water and corresponding diets. Data was collected real-time and analyzed as recommended by the manufacturers. The presented data were collected after 4 days of accommodation to the new environment. Energy expenditure and respiratory exchange ratio (RER) were measured by an indirect gas calorimetry module recording O2 consumption and CO2 production Locomotor and ambulatory activity was recorded as beam breaks converted into distance/velocity, measuring activity in three dimensions and analyzed in the metabolic cage using custom software. Data analysis and statistics were conducted with CalR software (https://www.calrapp.org).

### Serum and tissue sample collection

Final blood serum and tissues are harvested at the end of the 4^th^ week for further analysis. Blood was collected first with cardiac puncture, then mice are euthanized for tissue collection. Collected tissues were clamped immediately on dry ice and stored in -80 before analysis.

### Elisa

Serum hormone levels (Insulin, Glucagon, FGF21, GDF15) were measured by ELISA kits. Serum samples were collected at the end of monitoring with cardiac puncture, then centrifuged for 10 min at 4 °C to collect supernatant. Samples were stored at -80 before use, ELISAs were performed based on the kit protocols (Insulin: Crystal Chem 90080; Glucagon: Crystal Chem 81518; FGF21: BioVendor RD291108200R; GDF15: Abcam ab216947).

### LC-MS sample preparation

10–20 mg of tissue sample is weighed and subject for metabolite extraction. Soluble metabolites extraction was done by adding -20 °C 40:40:20 methanol: acetonitrile: water buffer at the ratio of 110 μl per 1 mg tissue. Samples were homogenized and extracted by bead homogenizer at 4°C, the resulting liquid samples were incubated at –80 °C overnight, thawed on ice and then centrifuged at 4 °C, 16,000g for 30 minutes. Supernatant was transferred to liquid chromatography–mass spectrometry (LC–MS) vials for analysis.

Hepatic glycogen extraction was done by adapting previous reported method^68^. Frozen weighed tissue was mixed with 2 M hydrochloric acid at 10 μl per mg tissue and then homogenized. The liquid was placed in water bath at 80°C for 2h before neutralizing with saturated ammonium bicarbonate at 12 μl per mg tissue. The resulting liquid was extracted by 88 μl 50:50 methanol: acetonitrile per mg of tissue, ending up in 10 mg tissue per 110 μl of samples. After overnight incubation at -80 °C, samples are centrifuged at 16,000g for 30 minutes before supernatant being analyzed. Glycogen amount was calculated by the acid -hydrolyzed glucose minus directly extracted glucose.

The saponified-fatty-acid extraction of tissue was adapted from previous research^67^. The extraction was done by adding 1 ml of 0.3 M KOH in 90:10 methanol: H_2_O buffer to the weighed tissue. After transferring the resulting mixture to a 4 ml glass vial, saponification was done at 80 °C in water bath for 1 hour. Then samples were cooled to room temperature, neutralized with 100 μL pure formic acid and extracted with 1 ml of hexane. The hexane layer was transferred to another glass vial, dried down under N2, and resuspended in 1:1 methanol: acetonitrile (100 μl per mg tissue; 40 μl per μl serum) for LC–MS analysis.

### LC-MS method

All LC-MS analysis was performed at the Proteomics Core at Rockefeller University. LC-MS analysis was conducted on a QExactive benchtop orbitrap mass spectrometer coupled to a Vanquish UPLC System (Thermo Fisher Scientific). External mass calibration was performed every 3 days using a standard calibration mixture. Polar and non-polar extracts were injected onto a ZIC-pHILIC (150 x 2.1mm) column and an Ascentis Express C18 (150 x 2.1mm column, respectively, using a previously described LC-MS method^69^.

Glucose/glycogen analysis was done with a newly developed method. Polar extracts (5 µl) were injected onto a ZIC-pHILIC 150 x 2.1mm (5μm particle size) column (EMD Millipore). Chromatographic separation was achieved using the following conditions mobile phase A consisted of 20 mM ammonium carbonate with 0.1% (v/v) ammonium hydroxide (adjusted to pH 9.3) and mobile phase B was acetonitrile. The column oven and autosampler tray were held at 40°C and 4°C, respectively. The chromatographic gradient was run at a flow rate of 0.150 mL/min as follows: 0-6 min: linear gradient from 70% to 55% B; 6-8 min: 55% to 40% B; 8-10 min: held at 40% B; 10-10.1 min: returned to 70% B; 10.1-16 min: equilibrated at 70% B. The mass spectrometer was operated in negative mode alternating between a full MS scan and a PRM scan. The spray voltage set to 3.0 kV, the heated capillary held at 275°C, the sheath gas flow was set to 40 units, and the auxiliary gas flow was set to 15 units. The MS1 data acquisition was acquired with the following parameters: scan range of 55-825 *m/z*, 140,000 resolution, 1 × 10^6^ AGC target, 80 ms injection time. The PRM scan was acquired with the following parameters: 17,500 resolution, 1 × 10^5^ AGC target, 50 ms injection time and stepped collision energy of 20, 30,40 units. The PRM scan was triggered by an inclusion listed targeting glucose and other isomers (C_6_H_12_O_6_) with the mass set to 179.0561 m/z.

### In vivo metabolic tracing

In-house catheterized mice are used for stable-isotope tracing to measure glucose turnover and glycogen sources. The method was adapted from previous research^33,67,68^. Glucose circulation rate was measured at ad lib fasted state (2pm-4:30pm), with 200mM in chow-fed mice and 50mM in KD-fed mice, at infusion rate of 0.1 µl/min/g. Glycogen tracing was done from 6pm to 10am (+1 day), with full access to food and water, and the infusate concentration are listed in the chart, with infusion rate of 0.1 µl/min/g.

| Tracer | Chow-fed | KD-fed |
| --- | --- | --- |
| U13C-glucose | 300mM | 50mM |
| U13C-alanine | 100mM | 100mM |
| U13C-glycerol | 100mM | 100M |

### Gfral-ablation

Gfral-cre (#036750) male mice were anesthetized with 3% and maintained at 1.5% isoflurane in oxygen, placed in a stereotax apparatus (Kofp Instruments) and kept warm with heat pads. 50nl of AAV5-hSyn-DIO-mCherry (2.2×10^13^, Addgene Cat # 50459-AAV5) or 50 nl AAV5-mCherry-flex-dTA (7×10^13^, Duke Viral Vector Core) viruses were injected into the area postrema (coordinates relative to the obex: AP: +0.2, ML: 0.0, DV: -0.5) using a glass pipette connected to a Nanoject (Drummond, Cat# 3-000-207). After finishing the injection, the glass capillaries were held for 10 minutes to allow proper diffusion of the virus before withdrawing it slowly. Animals were allowed to recover and express virus for 3 before diet change.

### Nnt overexpression

Nnt gene was synthesized by Twist Bioscience and cloned to a TBG promoter-containing AAV vector. Both Nnt-containing and control plasmid are sent to viraltools at Janelia farm to make AAVDJ/8 virus. Around 1e+12 titers of virus was delivered retro-orbitally to the mice and after 3-weeks, dietary challenge started.

### Immunoblotting

Tissues were homogenized using 2-ml homogenizer prior to lysis in triton lysis buffer (50 mM Tris–HCl, pH 7.5, 1 mM EDTA, 1% (v/v) Triton X-100, 150 mM NaCl) supplemented with protease inhibitor cocktail (EMD Millipore, 539134). Each lysate was sonicated, after centrifugation at 20,000 g at 4°C for 10 min, supernatants were collected. Sample protein concentrations were determined by using Pierce BCA Protein Assay Kit (Thermo Scientific) with bovine serum albumin as a protein standard. Samples were resolved on 10-20% SDS-PAGE gels (Invitrogen) and analyzed by standard immunoblotting protocol. Briefly, membranes were incubated with primary antibodies Nnt (ProteintecH, 13442-2-AP, 1:1000), GAPDH (GeneTex, GTX627408, 1:6000) in 5% (w/v) nonfat milk in TBST at 4°C overnight with shaking. Membranes were washed three times in TBST. Membranes were incubated with secondary antibody for 1 hour at room temperature with shaking, including α-mouse IgG-HRP linked (Cell Signaling, 7076) and α-rabbit IgG-HRP linked (Cell Signaling, 7074), used at 1: 5000 dilutions. Membranes were washed again three times in TBST before visualization with ECL substrate.

### Hepatic RNA sequence and data analysis

Frozen liver samples weighed 10-20mg and MRNA extraction was done with Qiagen RNeasy mini kit (#74106) following the protocol. RNAseq was done with the genetics core at Rockefeller University. For ad lib fasted state RNAseq, we collaborated with BGI. A certain amount of total RNA samples was taken and used oligo dT beads to enrich mRNA with poly A tail. After mRNA molecules were fragmented into small pieces, they were synthesized into first strand cDNA using random primers. The second strand cDNA was synthesized with dUTP instead of dTTP. Then, the synthesized cDNA was subjected to end-repair and 3’ adenylated. Adaptors were ligated to the ends of these 3’ adenylated cDNA fragments. Afterwards, we digested the U-labeled second-strand template with Uracil-DNA-Glycosylase (UDG) and perform PCR amplification. Library quality control and circularization was performed before sequencing. The library was then amplified to make DNA nanoball (DNB) and then sequenced on DNBSEQ (DNBSEQ Technology) platform.

For refed status RNAseq, we collaborated with the genetics core at Rockefeller University. 100 ng of total RNA was used to generate RNA-Seq libraries using Illumina stranded mRNA prep kit (Cat# 20040534) following manufacturer’s protocol. Libraries prepared with unique barcodes were pooled at equal molar ratios. The pool was denatured and sequenced on Illumina NextSeq 500 sequencer using high output V2 reagents and NextSeq Control Software v1.4 to generate 75 bp single reads, following manufacture’s protocol (Cat# 15048776 Rev.E)

For B6NJ and B6J comparison, we collaborated with the genetics core at Rockefeller University. 100 ng of total RNA was used to generate RNA-Seq libraries using Illumina stranded mRNA prep kit (Cat#20040534) following manufacturer’s protocol. Libraries prepared with unique barcodes were pooled at equal molar ratios. The pool was denatured and sequenced on Element AVITI sequencer using Cloudbreak FS reagents to generate 2 x 75 bp paired end reads, following manufactures protocol (Document # MA-00008 Rev. K).

Counts matrices were generated from the raw fastqs by pseudoalignment against either mm10 or hg38 using salmon^70^, and the resulting tables were loaded into R with tximport. Samples were normalized and variance-stabilized and differentially expressed genes were called through the limma-voom pipeline^71^. The logFC and p-values computed by limma-voom for all genes were used as inputs for pathway analysis using the fGSEA implementation. Secreted proteins were identified from the Secreted Protein Database (SEPDB)^72^, filtering only to those annotated with an SPB class of RANK0.

Metabolic inference was done with the following method: normalized and variance-stabilized counts for each sample were exported in TSV format and used as the input for Compass^29^ against either the human or the mouse reference. The resulting penalties were converted to flux estimates by taking the negative logarithm of one plus the penalty, and fluxes were then min-fixed and Z-scored. The effect size for the change in flux for each reaction was calculated as the Cohen’s d, or the mean divided by the standard deviation. These effect sizes were used for downstream visualization.

### Pharmacological administration

N-Acetyl-L-cysteine, BioXtra, ≥99% (TLC) (A8199-10G) was reconstituted in PBS (adjust PH with NaOH to neutral) and injected intraperitoneally (i.p.) daily at 250mg/kg body weight while fed with different ketogenic diets. Isrib (Mechem Express #HY12495) was first dissolved in DMSO at 5 mg/ml, then diluted in 20% PEG 300 and 2% Tween 80 (in PBS) to a final concentration of 0.25 mg/ml. Mice are given Isrib at 2.5mg/kg every other day via i.p. injections, which were all done at indicated concentrations using insulin syringes (Beckton Dickinson, Cat# 324911). Control groups are given vehicle i.p. daily.

## Acknowledgements

We thank all members of the laboratory of J.M.F. for discussions and support, Dr. Connie Zhao from the Genomics Resource Center for technical assistance with RNA-seq experiments. Dr. Paul Cohen for the metabolic cage experiments. This work was supported by the JPB Foundation and the Howard Hughes Medical Institute (to J.M.F.), the Helen Hay Whitney Foundation Postdoctoral Fellowship (to Z.Z), and the Maurice R. and Corinne P. Greenberg Center for the Advancement of Translational Research Pilot Project Grant (to Z.Z). Illustration was created with Biorender (https://biorender.com).

## Author contributions

Z.Z conceived the study, designed, performed and analyzed all experiments with input from all authors. A.M.A performed GFRAL-neuron ablation experiment. S.L performed virus delivery and western blot in the NNT overexpression. A.M. performed all bioinformatic analysis. J.S. helped maintain mouse lines and genotyping. H. A., M.I. and H.M. helped design MS methods and run samples. K.B provides input in experimental design. Z.Z and J.M.F. wrote the manuscript.

---

**Supplementary figure 1.**
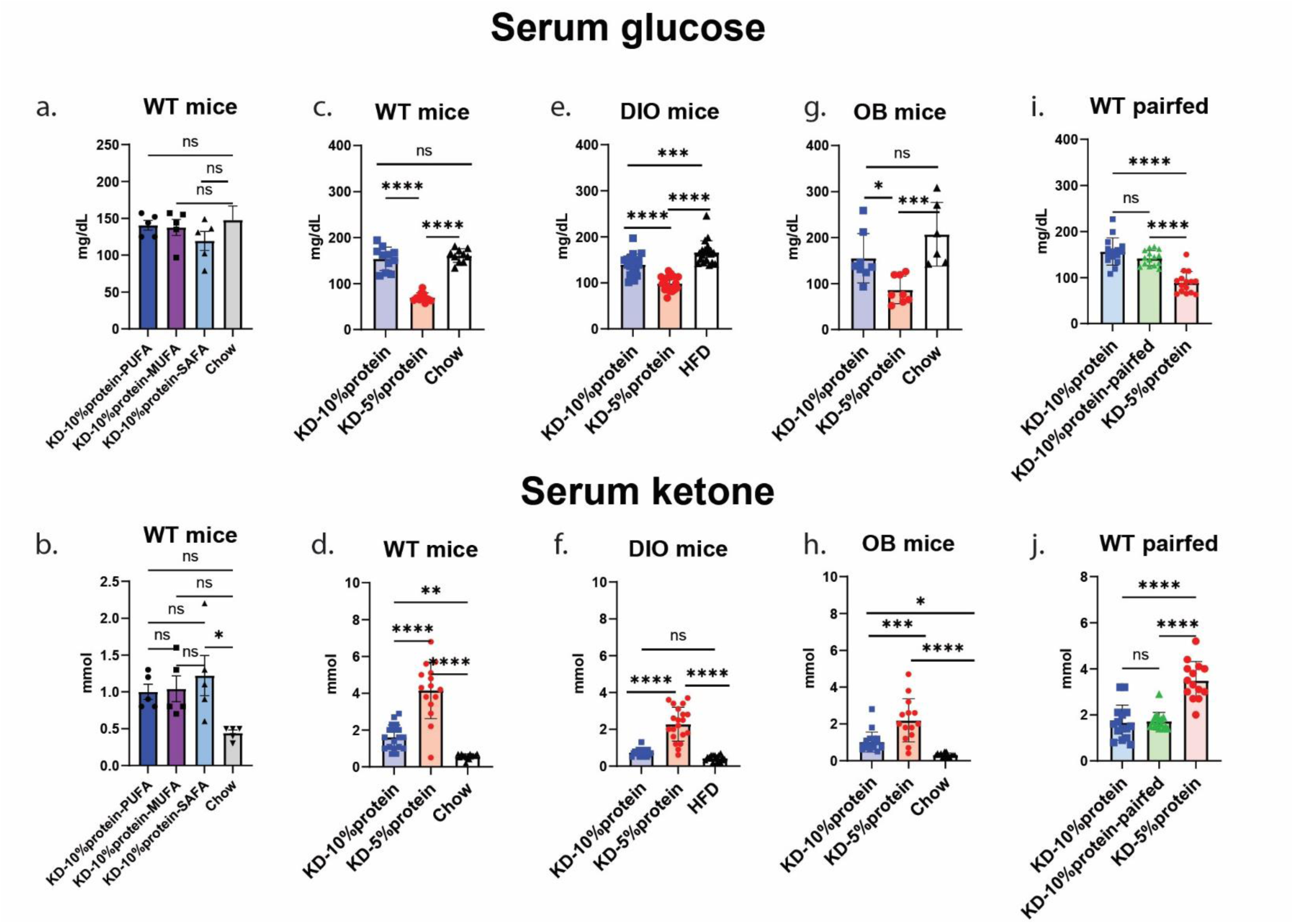
Protein Content Dictates glucose and ketone levels in B6J mice. a. Serum glucose level of WT B6J mice under KD-10%P rich in PUFA, MUFA, SAFA and chow. b. Serum ketone level of WT B6J mice under KD-10%P rich in PUFA, MUFA, SAFA and chow. c-d. As in a-b, for WT mice under KD-10%P or KD-5%P rich in PUFA and chow. e-f. As in a-b, for DIO mice (control group is HFD). g-h. As in a-b, for OB mice. i-j. As in a-d, for WT B6J pair-feeding mice. (n = 5 mice for all WT test, n=4 for all DIO test, n=4 for OB on KD and n=3 for OB on chow. All tests are done by testing 2 individual days of the same group of mice, besides panel a and b, which tested one time. Data are presented as mean ± s.e.m. NS, not significant; *P < 0.05; **P < 0.01; ***P < 0.001; ****P < 0.0001.)

**Supplementary figure 2.**
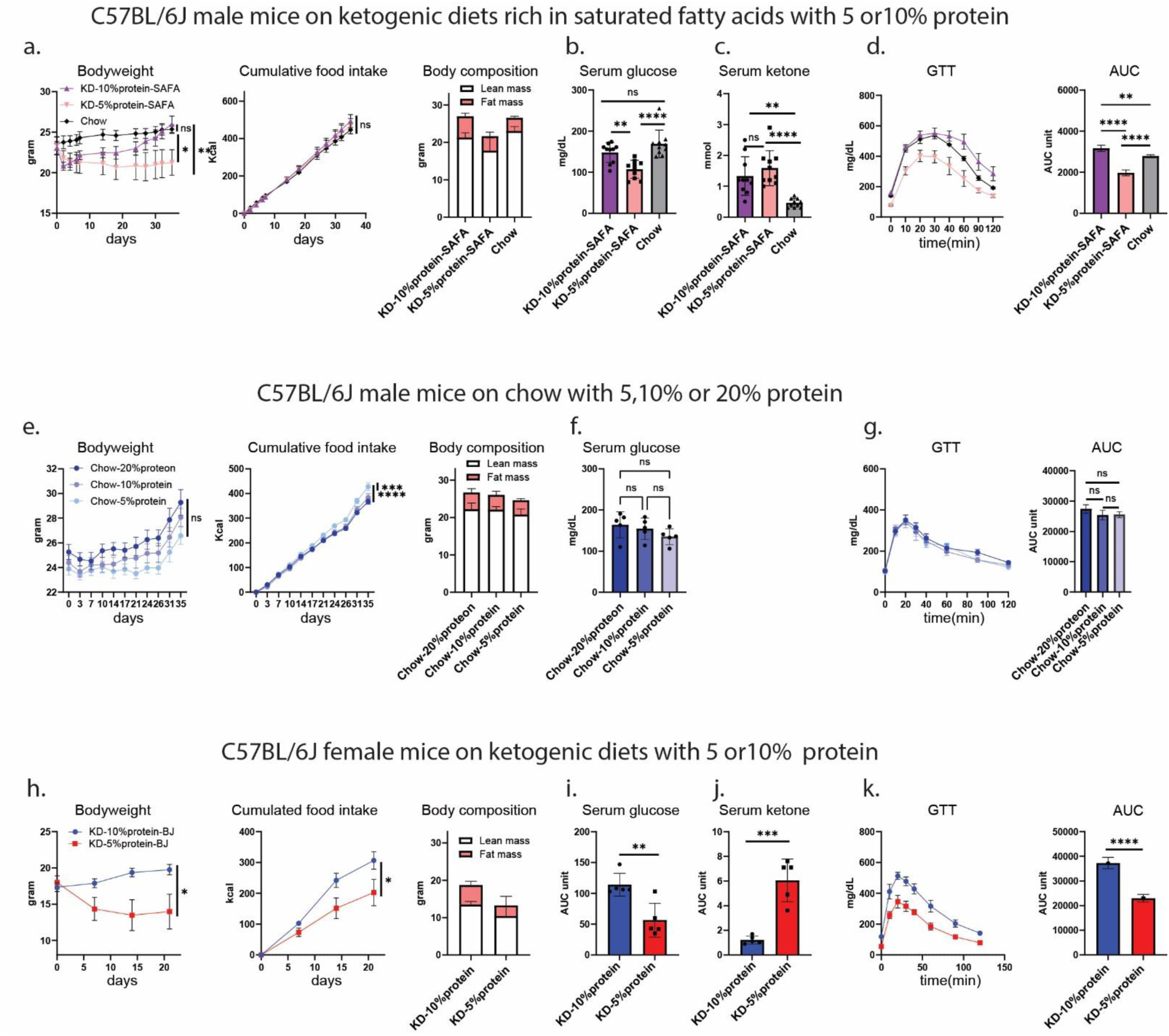
Phenotypes of B6J male and female mice on other custom diets. a. Body weight, cumulative calorie intake, and body composition of B6J male mice on KD-10%P or KD-5%P rich in SAFA. b. Serum glucose of B6J male mice on KD-10%P or KD-5%P rich in SAFA. c. Serum ketone of B6J male mice on KD-10%P or KD-5%P rich in SAFA. d. Glucose tolerance test (2mg/gram body weight) and calculated area under the curve of B6J male mice on KD-10%P or KD-5%P rich in SAFA. e, f. As in a-b, for B6J male mice on chow with 5%,10% or 20% protein. g. Glucose tolerance test (2mg/gram body weight) and calculated area under the curve of B6J male mice on chow with 5%,10% or 20% protein. h-k. As in a-f, for B6J female mice on KD-10%P or KD-5%P. (n = 5 mice for all tests. Data are presented as mean ± s.e.m. NS, not significant; *P < 0.05; **P < 0.01; ***P < 0.001; ****P < 0.0001.)

**Supplementary figure 3.**
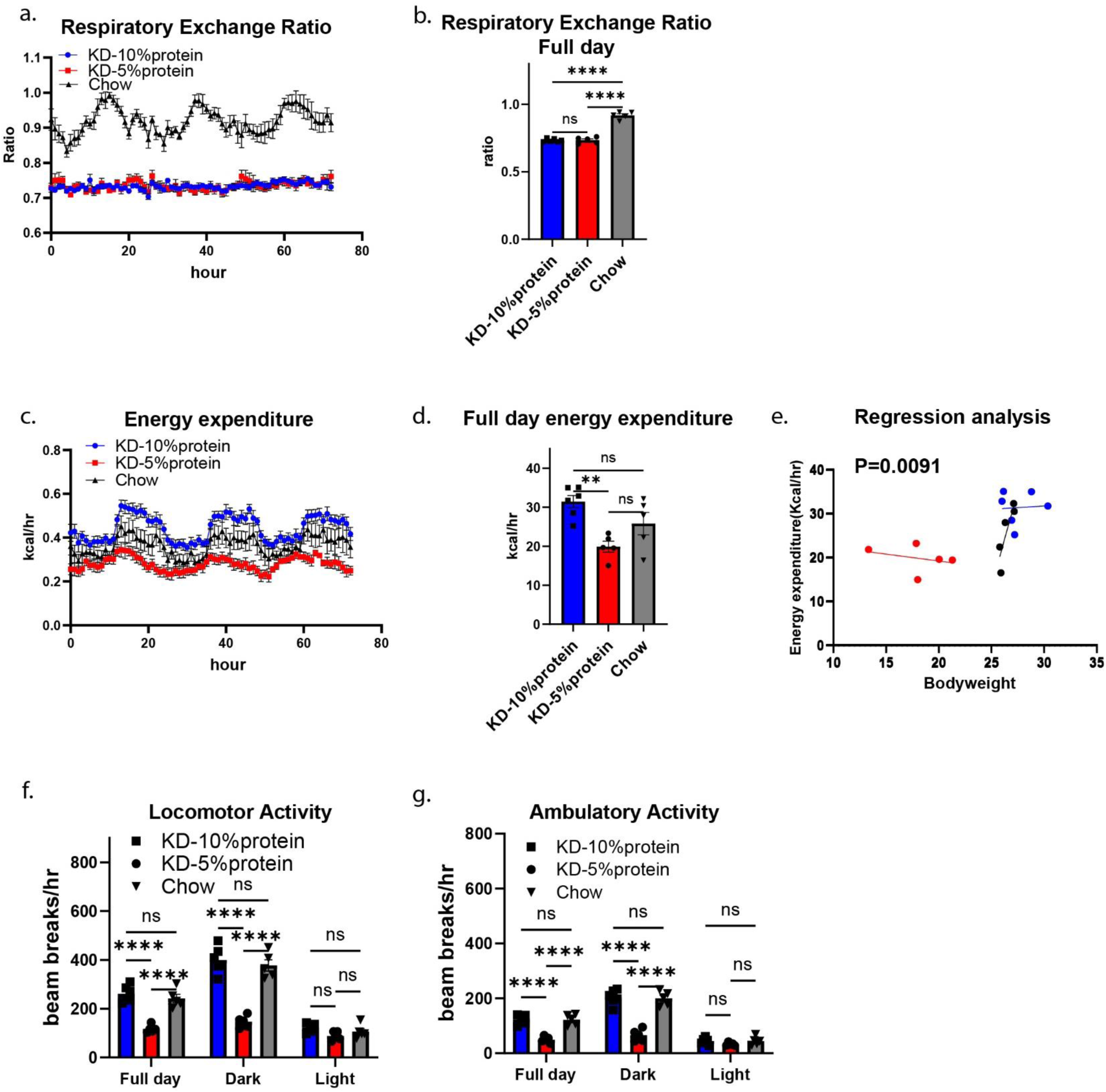
KD with 5% or 10% protein alters energy expenditure and activities in B6J mice. a. Respiratory exchange ratio of B6J mice under KD-5%P, KD-10%P or chow. b. Full day averaged respiratory exchange ratio of B6J mice under KD-5%P, KD-10%P or chow. c. Energy expenditure of B6J mice under KD-5%P, KD-10%P or chow. d. Full day averaged energy expenditure of B6J mice under KD-5%P, KD-10%P or chow. e. Energy expenditure regression analysis of B6J mice under KD-5%P, KD-10%P or chow. f. Averaged locomotor activity in different time phase of B6J mice under KD-5%P, KD-10%P or chow. g. Averaged ambulatory activity in different time phase of B6J mice under KD-5%P, KD-10%P or chow. (n = 6 mice for KD-10%P group and n=5 for KD-5%P and chow groups. Data are presented as mean ± s.e.m. NS, not significant; *P < 0.05; **P < 0.01; ***P < 0.001; ****P < 0.0001.)

**Supplementary figure 4.**
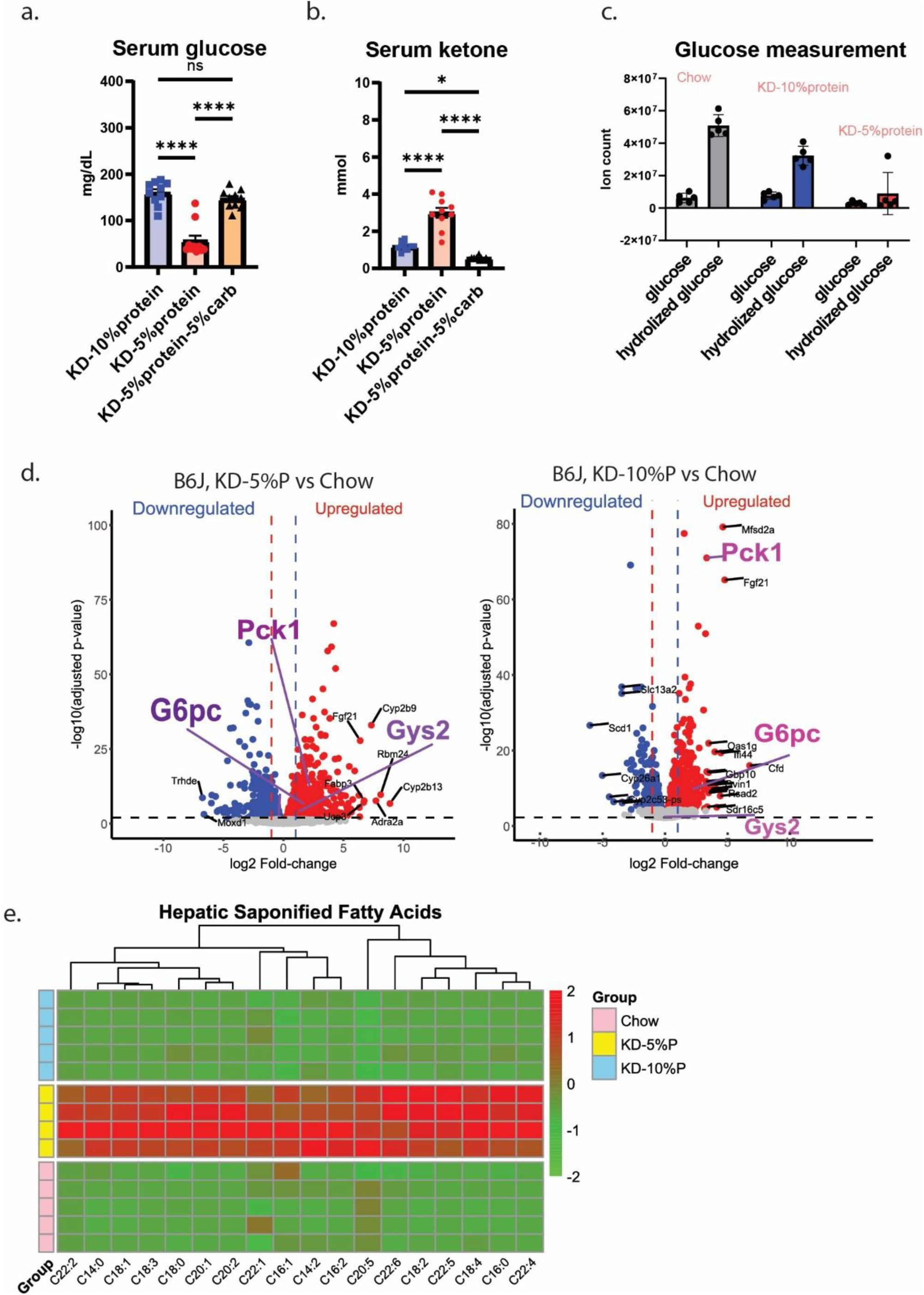
KD with different composition remodeled metabolism. a. Serum glucose of WT B6J mice fed with KD-5%P, KD-10%P and KD-5%P-5%C. b. Serum ketone of WT B6J mice fed with KD-5%P, KD-10%P and KD-5%P-5%C. c. Raw data of hepatic glucose and hydrolyzed glucose levels of WT B6J mice fed with KD-5%P, KD-10%P and chow d. Hepatic transcription of KD-10%P-fed and KD-5%P-fed mice versus chow-fed mice. e. Hepatic saponified fatty acids profile in mice fed with KD-5%P, KD-10%P and chow. (n = 5 mice for each test. All glucose/ketone are done by testing 2 individual days of the same group of mice. Data are presented as mean ± s.e.m. NS, not significant; *P < 0.05; **P < 0.01; ***P < 0.001; ****P < 0.0001.)

**Supplementary figure 5.**
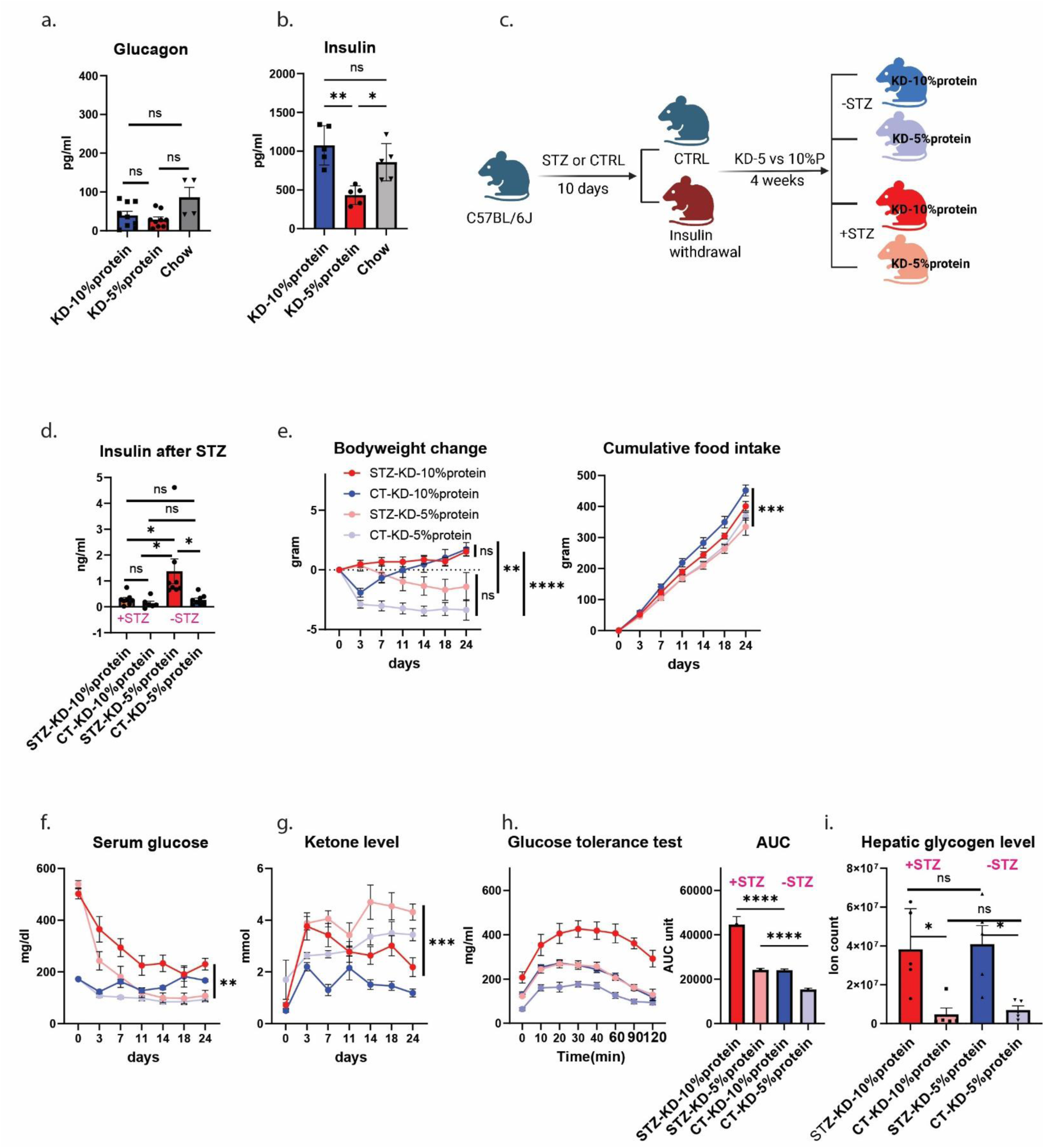
The glucose homeostasis maintenance under KD is insulin independent. a. Circulating glucagon levels of WT B6J mice under KD-5%P, KD-10%P or chow. b. Circulating insulin levels of WT B6J mice under KD-5%P, KD-10%P or chow. c. The schematic of STZ-induced insulin withdrawal in WT B6J mice and feed with KD-5%P or KD-10%P. d. Circulating insulin levels of WT B6J mice treated with or without STZ under KD-5%P, KD-10%P. e. Body weight change and cumulative calorie intake (Kcal) of WT B6J mice treated with or without STZ under KD-5%P, KD-10%P. f. Serum glucose levels of WT B6J mice treated with or without STZ under KD-5%P, KD-10%P. g. Serum ketone levels of WT B6J mice treated with or without STZ under KD-5%P, KD-10%P. h. Glucose tolerance test (1mg glucose/kg body weight) and calculated area under the curve of WT B6J mice treated with or without STZ under KD-5%P, KD-10%P. i. Hepatic glycogen levels of WT B6J mice treated with or without STZ under KD-5%P, KD-10%P. (n = 8 mice for all groups except CT-KD-10%protein(n=6). Data are presented as mean ± s.e.m. NS, not significant; *P < 0.05; **P < 0.01; ***P < 0.001; ****P < 0.0001.)

**Supplementary figure 6.**
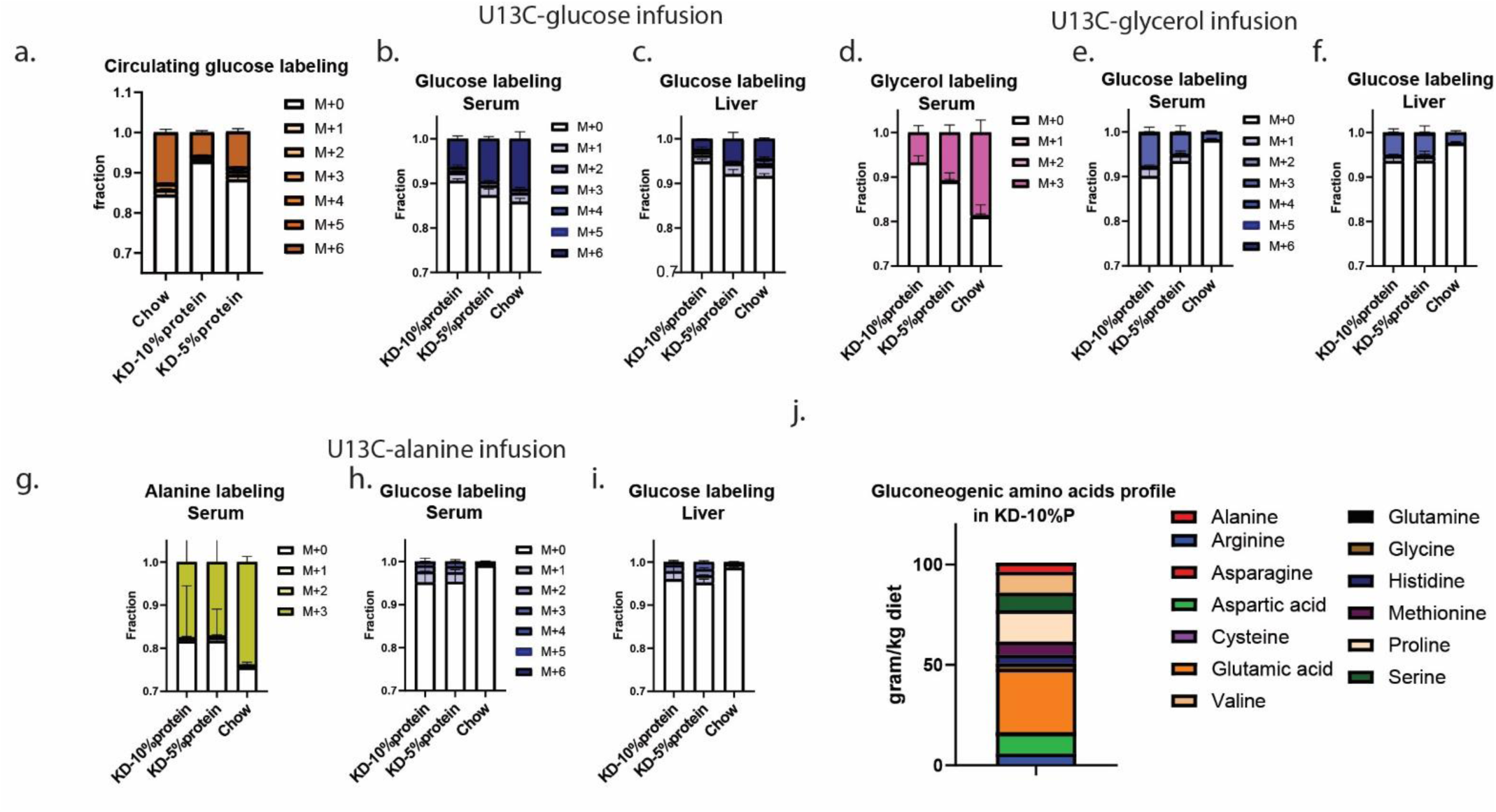
Raw tracing data and dietary amino acids profile of mice fed with KDs. a. Raw circulating glucose labeling after 2.5h U^13^C-glucose infusion at fasted state in B6J mice under KD-5%P, KD-10%P or chow. b. Raw circulating glucose labeling after 16h U^13^C-glucose infusion at fed state in B6J mice under KD-5%P, KD-10%P or chow. c. As in b, for glucose labeling in liver d. Raw circulating glycerol labeling after 16h U^13^C-glycerol infusion at fed state in B6J mice under KD-5%P, KD-10%P or chow. e. As in d, for circulating glucose labeling. f. As in d, for glucose labeling in liver. g. Raw circulating alanine labeling after 16h U^13^C-glycerol infusion at fed state in B6J mice under KD-5%P, KD-10%P or chow. h. As in g, for circulating glucose labeling. i. As in g, for glucose labeling in liver. j. Dietary amino acids profile in KD-10%P. (Raw data are listed in source file) (n = 5 mice for glucose turnover measurement. For alanine tracing, n=3 for KD-5%P, n=3 for KD-10%P, n=5 for chow; for glycerol tracing n=4 for KD-5%P, n=4 for KD-10%P, n=4 for chow; for glucose tracing n=2 for KD-5%P, n=4 for KD-10%P, n=3 for chow. Data are presented as mean ± s.e.m. NS, not significant; *P < 0.05; **P < 0.01; ***P < 0.001; ****P < 0.0001.)

**Supplementary figure 7.**
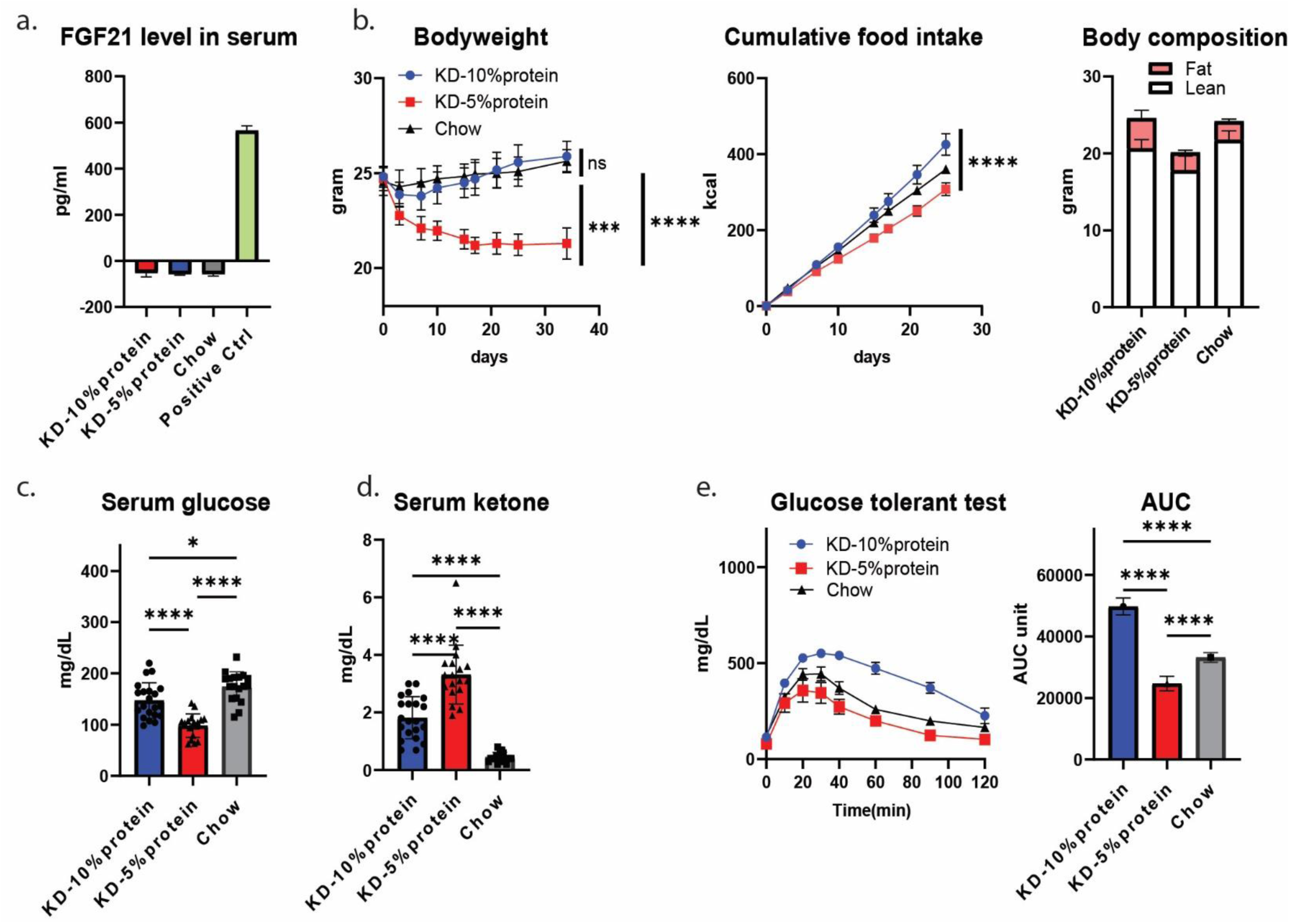
The different response to KD-5%P versus KD-10%P is FGF21 independent. a. Circulating FGF21 levels of FGF21 whole-body knockout mice under KD-5%P, KD-10%P or chow. b. Body weight, cumulative calorie intake, and body composition of FGF21 whole-body knockout mice under KD-5%P, KD-10%P or chow. c. Serum glucose levels of FGF21 whole-body knockout mice under KD-5%P, KD-10%P or chow. d. Serum ketone levels of FGF21 whole-body knockout mice under KD-5%P, KD-10%P or chow. e. Glucose tolerance test (2mg glucose/kg body weight) and calculated area under the curve of FGF21 whole-body knockout mice under KD-5%P, KD-10%P or chow. (n = 7 for KD-10%P group, n=6 for KD-5%P and chow groups. All glucose/ketone are done by testing 2 individual days of the same group of mice. Data are presented as mean ± s.e.m. NS, not significant; *P < 0.05; **P < 0.01; ***P < 0.001; ****P < 0.0001.)

**Supplementary figure 8.**
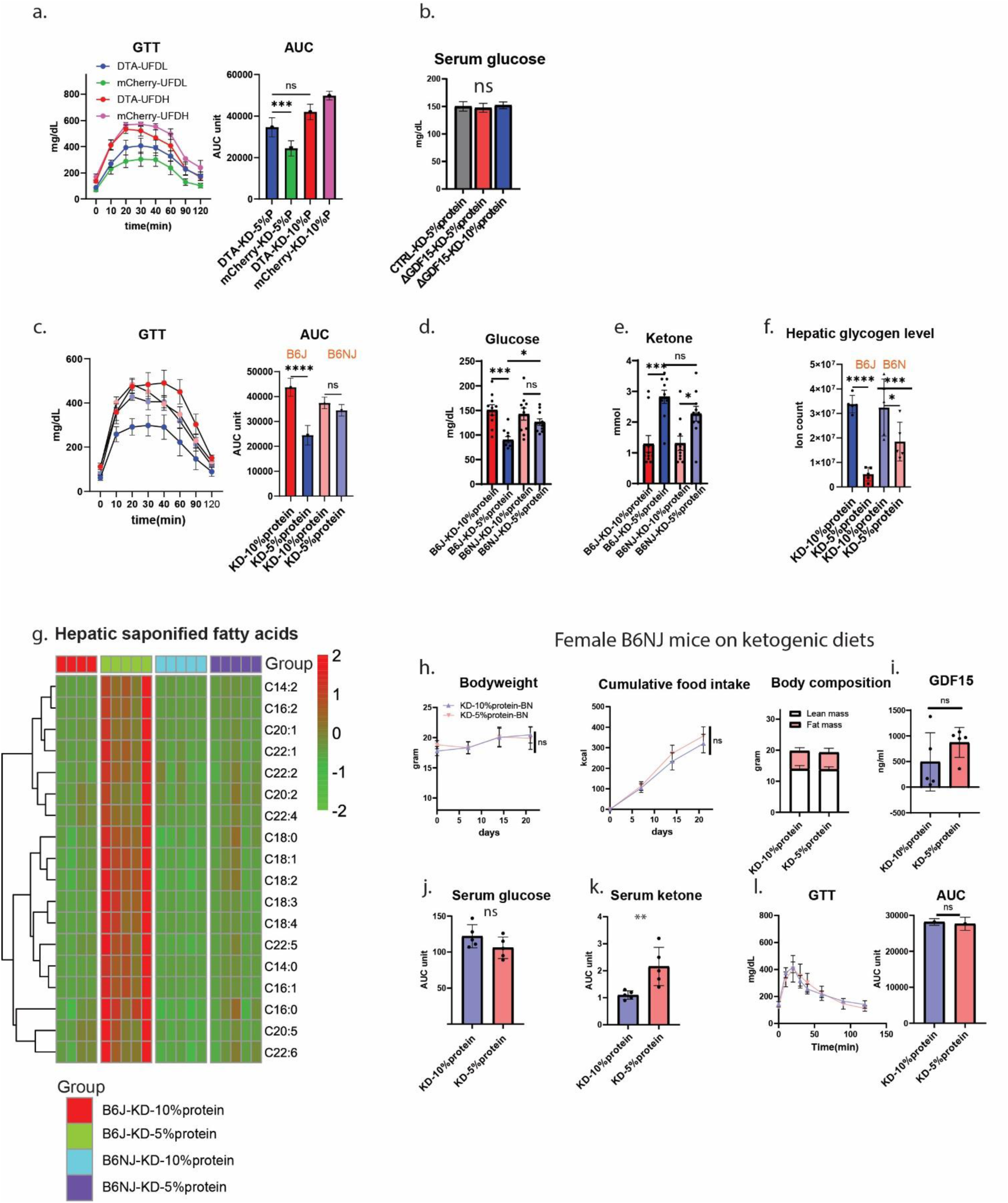
Male and female B6NJ mice response similarly towards KD-5%P or KD-10%P. a. Glucose tolerance test (2mg glucose/kg body weight) and calculated area under the curve of mice ablating GFRAL-neurons in Area Postrema and control mice under KD-5%P or KD-10%P. b. Serum glucose levels of B6NJ-Gdf15 whole-body knockout mice under KD-5%P or KD-10%P. c. Glucose tolerance test (2mg glucose/kg body weight) and calculated area under the curve of WT B6J or B6NJ mice under KD-5%P or KD-10%P. d. Serum glucose levels of WT B6J or B6NJ mice under KD-5%P or KD-10%P. e. Serum ketone levels of WT B6J or B6NJ mice under KD-5%P or KD-10%P. f. Hepatic glycogen levels of WT B6J or B6NJ mice under KD-5%P or KD-10%P. g. Hepatic saponified fatty acids profiles of WT B6J or B6NJ mice under KD-5%P or KD-10%P. h. Body weight, cumulative calorie intake (Kcal), and body composition of female WT B6NJ mice under KD-5%P or KD-10%P. i. Circulating GDF15 levels of female WT B6NJ mice under KD-5%P or KD-10%P. j. Serum glucose levels of female WT B6NJ mice under KD-5%P or KD-10%P. k. Serum ketone levels of female WT B6NJ mice under KD-5%P or KD-10%P. l. Glucose tolerance test (2mg glucose/kg body weight) and calculated area under the curve of female WT B6NJ mice under KD-5%P or KD-10%P. (n = 5 for all groups. All glucose/ketone are done by testing 2 individual days of the same group of mice. Data are presented as mean ± s.e.m. NS, not significant; *P < 0.05; **P < 0.01; ***P < 0.001; ****P < 0.0001.)

**Supplementary figure 9.**
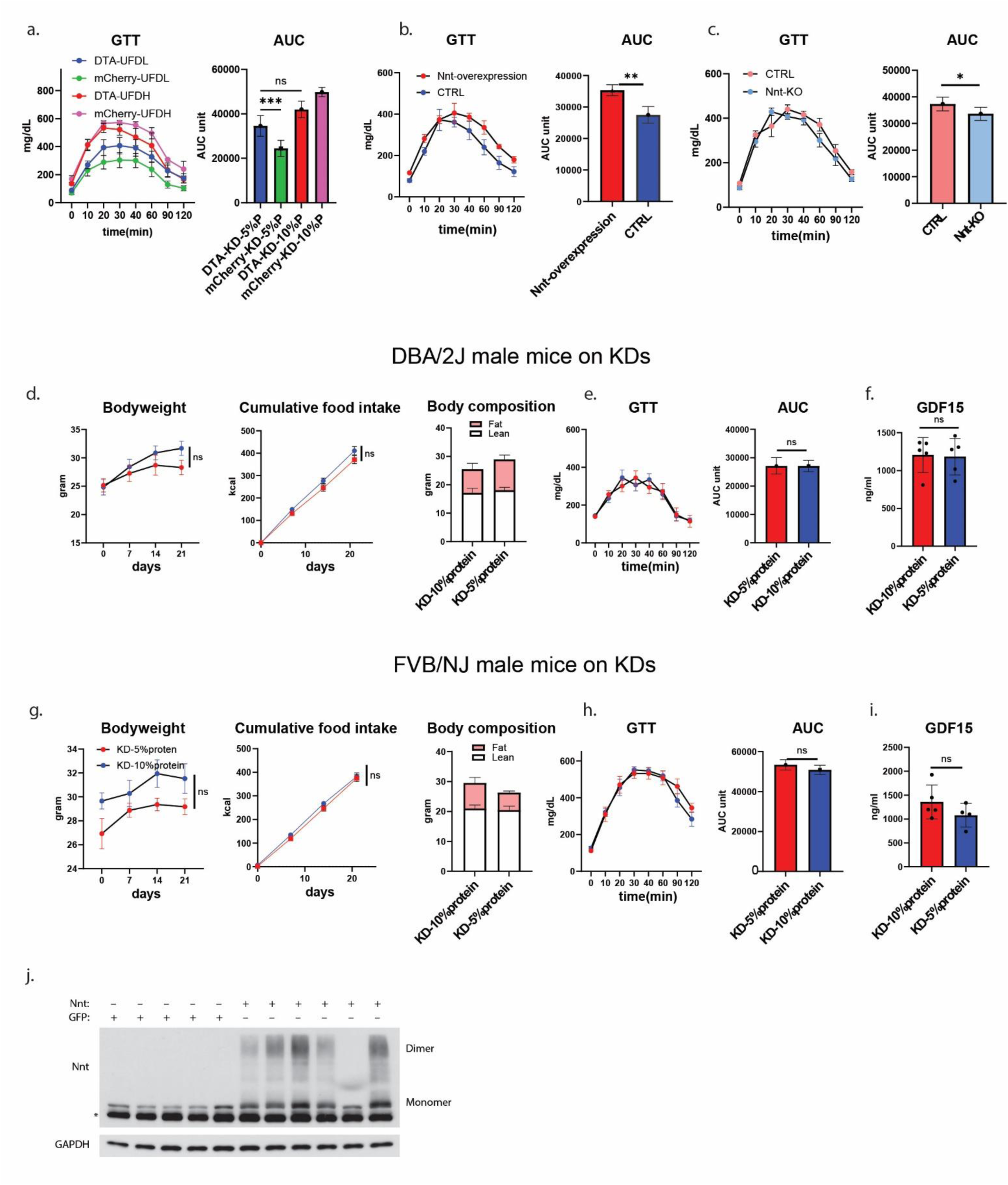
Other strains with functional Nnt show similar responses towards KD-5%P and KD-10%, while genetically manipulation of Nnt regulates the phenotypes. a. Glucose tolerance test (2mg glucose/kg body weight) and calculated area under the curve of Gfral-ablation and control mice under KD-5%P or KD-10%P. b. Glucose tolerance test (2mg glucose/kg body weight) and calculated area under the curve of B6NJ-Nnt whole-body knockout mice under KD-5%P or KD-10%P. c. Glucose tolerance test (2mg glucose/kg body weight) and calculated area under the curve of B6J mice with NNT re-expression under KD-5%P or KD-10%P. d. Body weight, cumulative calorie intake (Kcal) and body composition of DBA/2J mice on KD-5%P or KD-10%P. e. Glucose tolerance test (2mg glucose/kg body weight) and calculated area under the curve of composition of DBA/2J mice under KD-5%P or KD-10%P. f. GDF15 levels of DBA/2J mice under KD-5%P or KD-10%P. g-i. As in d-f, for FVB/NJ mice. j. Western blot of Nnt in the liver of control or overexpression mice. (n = 5 for each group in DBA/2J, n=4 for each group in FVB/2J. All glucose/ketone are done by testing 2 individual days of the same group of mice. Data are presented as mean ± s.e.m. NS, not significant; *P < 0.05; **P < 0.01; ***P < 0.001; ****P < 0.0001.)

**Supplementary figure 10.**
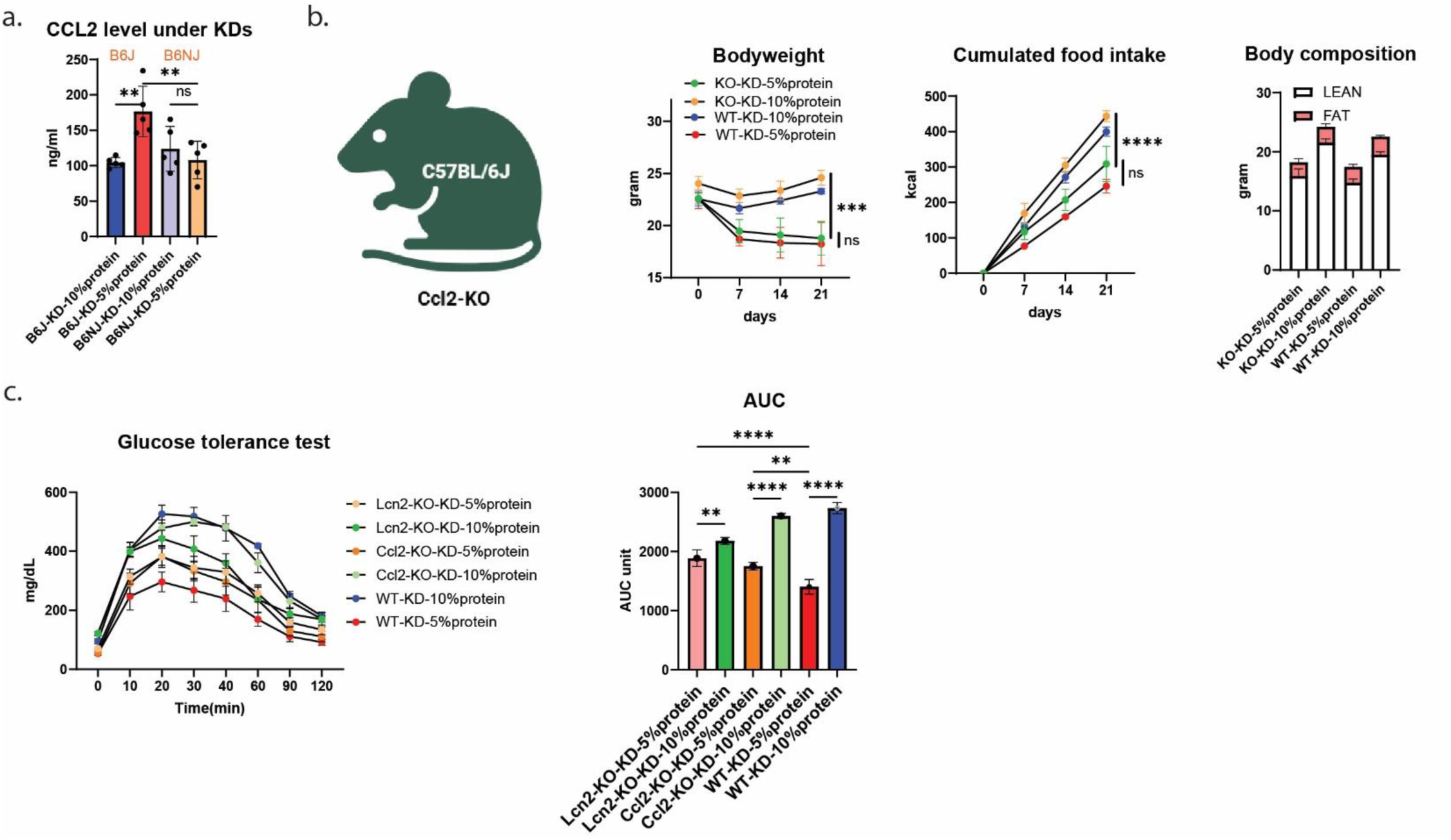
The different response to KD-5%P versus KD-10%P is CCL2 independent. a. Serum CCL2 levels of WT B6J or B6NJ under KD-5%P or KD-10%P. b. Body weight, cumulative calorie intake (Kcal) and body composition of B6J-CCL2 whole-body knockout and WT B6J mice on KD-5%P or KD-10%P. c. Glucose tolerance test (2mg glucose/kg body weight) and calculated area under the curve of B6J-LCN2, B6J-CCL2 whole-body knockout and WT B6J mice on KD-5%P or KD-10%P. (n = 5 for each group in panel a. For B6J-CCL2 KO experiment, n=5 for KO under KD-5%P group and n=4 for all other groups. Data are presented as mean ± s.e.m. NS, not significant; *P < 0.05; **P < 0.01; ***P < 0.001; ****P < 0.0001.)

**Supplementary figure 11.**
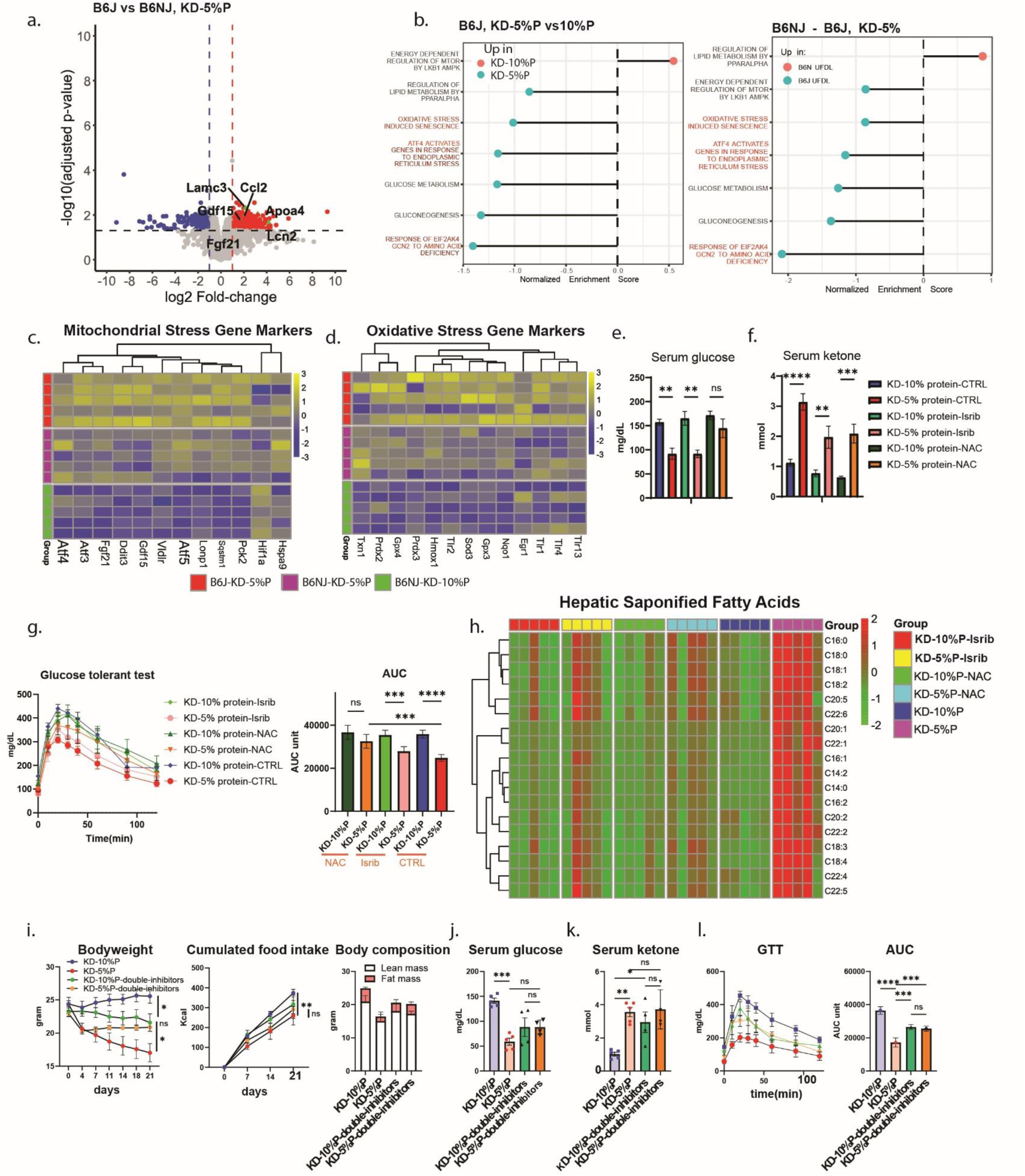
Stress pathway plays a role in the strain-specific response to KD-5%P, antioxidant can reverse it. a. Volcano plot of hepatic transcriptome of B6J versus B6NJ under KD-5%P. b. Pathway analysis of hepatic transcriptome of B6J under KD-5%P versus KD-10%P and B6J versus B6NJ under KD-5%P. c. Mitochondrial stress gene markers of hepatic transcriptome of B6J B6NJ under KD-5%P and B6NJ under KD-10%P. d. Oxidative stress gene markers of hepatic transcriptome of B6J B6NJ under KD-5%P and B6NJ under KD-10%P. e. Serum glucose of B6J mice treated with Isrib, NAC or Vehicle under KD-5%P or KD-10%P. f. Serum ketone of B6J mice treated with Isrib, NAC or Vehicle under KD-5%P or KD-10%P. g. Glucose tolerance test (2mg glucose/kg body weight) and calculated area under the curve of B6J mice treated with Isrib, NAC or Vehicle under KD-5%P or KD-10%P h. Hepatic fatty acids profile of B6J mice treated with Isrib, NAC or Vehicle under KD-5%P or KD-10%P. i. Body weight, cumulative calories intake (Kcal) and body composition of control B6J and Isrib-NAC double-treated mice under KD-5%P or KD-10%P. j. Serum glucose of control B6J and Isrib -NAC double-treated mice under KD-5%P or KD-10%P. k. Serum ketone of control B6J and Isrib -NAC double-treated mice under KD-5%P or KD-10%P. l. Glucose tolerance test (2mg glucose/kg body weight) and calculated area under the curve of control B6J and Isrib -NAC double-treated mice under KD-5%P or KD-10%P. (n = 5 for each group in a-g. n=5 for both control groups and n=4 for double inhibitor experiments. All glucose/ketone are done by testing 2 individual days of the same group of mice. Data are presented as mean ± s.e.m. NS, not significant; *P < 0.05; **P < 0.01; ***P < 0.001; ****P < 0.0001.)

